# Optimizing cytochrome P450 activity for quillaic acid biosynthesis in *Saccharomyces cerevisiae*

**DOI:** 10.1101/2025.11.09.685466

**Authors:** Yutong Han, Laura Coe, James J. De Voss, Birgitta E. Ebert

**Affiliations:** Australian Institute for Bioengineering and Nanotechnology, The University of Queensland, St Lucia, QLD 4072, Australia; School of Chemistry and Molecular Biosciences, The University of Queensland, St. Lucia, QLD 4072, Australia; Food and Beverage Accelerator (FaBA), The University of Queensland, Brisbane, Queensland, Australia

**Keywords:** *Saccharomyces cerevisiae*, cytochrome P450, triterpenoid, plant natural product, mevalonate pathway, GAL promoters, redox cofactor supply, synthetic biology

## Abstract

QS-21, a saponin extract from the Chilean tree *Quillaja saponaria*, is gaining popularity as a potent vaccine adjuvant, but its production is constrained due to its low natural abundance and complex chemical structure of the triterpenoid saponins, which hinder large-scale production through plant extraction or chemical synthesis. Microbial biosynthesis presents a promising alternative, with *Saccharomyces cerevisiae* emerging as a desirable host for triterpenoid production. A key step toward microbial QS-21 synthesis is the efficient biosynthesis of its aglycone core, the triterpenoid quillaic acid. However, its efficient production remains limited by challenges of cytochrome P450 enzyme (CYP450s) activity, including cofactor availability and electron transfer efficiency.

To address this limitation, we applied a multi-faceted metabolic engineering strategy to optimise CYP450 activity, including CYP450 expression and cytochrome P450 reductase (CPR) selection. Additionally, aligning CYP450 expression with the ethanol phase, enhanced the metabolic flux toward quillaic acid synthesis, leading to an 85-fold increase in titre. Together, these strategies led to a quillaic acid titre of 385 ± 14 mg/L in flask fermentation. Fed-batch bioreactor fermentations increased quillaic acid titre to 471 ± 20 mg/L and significantly increased the selectivity for QA from 32.6% to 65.1% of the total triterpenoids produced. These findings demonstrate the effectiveness of enhancing CYP450 activity through targeted strategies and reveal potential bottlenecks in CYP450 expression, providing valuable insights for future optimization of triterpenoid production in yeast.

## 1. Introduction

The rapid advancement in the development of microbial cell factories allows to explore the recombinant production of complex natural products and create new artificial products (Liu et al., 2022). Compared with traditional extraction from plants and chemical synthesis, microbial synthesis offers various advantages such as short growth cycle, simple operation, environmental friendliness, and controllable large-scale fermentation (Srinivasan and Smolke, 2020). Within this space, triterpenoids, a class of structurally diverse metabolites widely found in plants, have attracted great attention for application in the pharmaceutical, food and cosmetics industries (Dinday and Ghosh, 2023). To date, the recombinant synthesis of several bioactive triterpenoids has been successfully implemented in various microbes (Guo et al., 2022; Guo et al., 2020). Among them, *Saccharomyces cerevisiae* stands out as a particularly desirable host due to its endogenous mevalonate (MVA) and sterol pathways providing the common triterpenoid precursor 2,3-oxidosqualene, the presence of endoplasmic reticulum (ER) to anchor membrane-associated plant enzymes, and the plethora of genetic tools for metabolic engineering (Brown et al., 2015; Kulagina et al., 2021; T. Liu et al., 2022; Nowrouzi et al., 2020; Qu et al., 2015). The biosynthesis pathway of triterpenoids in yeast can be established by cyclisation of 2,3-oxidosqualene and site-specific oxidations catalysed by cytochrome P450 monooxygenases (CYP450s). The effectiveness of heterologous CYP450 expression is crucial for triterpenoid biosynthesis (Jiang et al., 2021) but still poses significant challenges, hindering the efficient production of triterpenoids in yeast (Zhou et al., 2021).

Cytochrome P450s (CYPs) are haem-thiolate monooxygenases that receive electrons from NADPH via cytochrome P450 reductase (CPR) to drive catalysis (Modi and Dawson, 2015; Quintanilha et al., 2017). Strategies to enhance heterologous CYP performance include increasing NADPH availability, selecting compatible CYP-CPR pairs and their ratio, and expanding the endoplasmic reticulum (ER) to better support membrane-associated enzymes. In yeast, screening plant CPR isoforms measurably shifted triterpenoid titres (Istiandari et al., 2021), and fine-tuning CPR relative to CYP improved CYP88D6-dependent 11-oxo-β-amyrin formation (Sun et al., 2023). Upregulation of *ICE2* promotes ER membrane biogenesis and stabilizes CPR, improving activity while maintaining ER homeostasis (Emmerstorfer et al., 2015; Papagiannidis et al., 2021). However, combining these interventions can introduce proteostatic and redox burdens; thus, gains are often incremental and context-dependent (Cha et al., 2022)

Ethanol has been described as preferred carbon source for triterpenoid production (Czarnotta et al., 2017). It directly supplies cytosolic acetyl-CoA, entry molecule of the MVA pathway, thus supporting higher metabolic flux toward triterpenoid biosynthesis (Sun et al., 2021) and increased NADPH availability to support CYP450 activity and other enzymes in the MVA pathway, such as the rate-limiting HMG-CoA reductase (HMGR) (Kim et al., 2018). A study by Zhu et al. used a glucose-sensing toggle switch to shift CYP450 expression to the ethanol phase, which significantly enhanced protopanaxadiol production in yeast (Zhu et al., 2022). These findings underscore the importance of matching CYP450 expression with optimal metabolic phases to improve production yields.

In this study, we investigated the impact of carbon source-dependent expression of recombinant plant CYP450s on quillaic acid biosynthesis in yeast. Quillaic acid is the triterpenoid core structure of the plant natural product QS-21 (Fig. 1b), one of the most potent vaccine adjuvants (Garçon and Van Mechelen, 2011; Jobe et al., 2022; Özverel et al., 2020; Ragupathi et al., 2011). Currently, the only approved source of QS-21 is the Chilean tree *Quillaja saponaria* (Wang et al., 2019), which makes supply limited and unstable due to destructive harvesting, long renewal times, and increasing protection concerns. Total and partial chemical synthesis of quillaic acid, and even more so QS-21, is not an economical option. Chemical synthesis of quillaic acid from the triterpenoid precursor oleanolic acid via C-H activation at C-16 and C-23 was reported in 14 steps, resulting in an overall yield of only 3.4%. Recent studies have demonstrated quillaic acid biosynthesis in yeast (Wang et al., 2024). For instance, Yang et al. achieved a titre of 258 mg/L by employing combinatorial engineering strategies (Yang et al., 2023), while a more recent study reported a titre of 2.23 g/L by optimizing the CYP450 network and spatial remodelling of enzymes (Wang et al., 2024). The recombinant production pathway was successfully extended to the two QS-21 isomers. Although both were obtained in only trace amounts (Liu et al., 2024), these achievements highlight the potential of QS-21 production in yeast.

**Fig 1.**
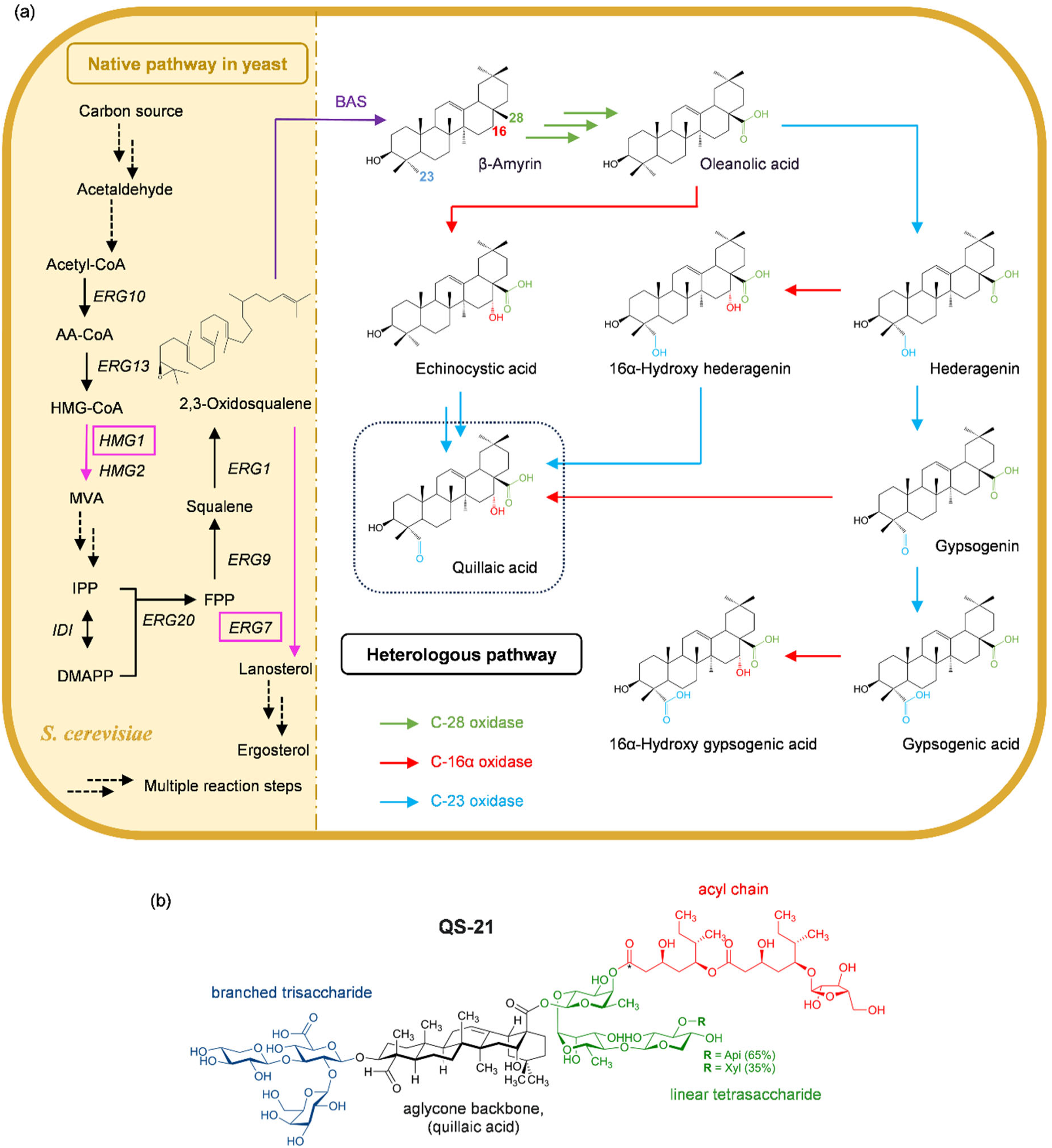
Recombinant biosynthesis of quillaic acid in yeast. (a) The triterpenoid precursor 2,3-oxidosqualene is derived from the common substrates glucose or ethanol via the native MVA pathway (left-hand side, highlighted in yellow). 2,3-oxidosqualene availability is commonly enhanced through deregulation of HMG-CoA reductase, the major limiting step of the MVA pathway, encoded by *HMG1* and *HMG2,* and downregulation of lanosterol synthase, the first committed step of the competing ergosterol biosynthesis, encoded by *ERG7*. 2,3-oxidosqualene is cyclised to β-amyrin, which is oxidised to quillaic acid through the activity of C-28, C-16 and C-23 oxidases. Due to the substrate promiscuity of CYP450 the exact sequence of oxidation steps is unknown and possible alternative pathways are indicated. (b) Chemical structure of QS-21; BAS, beta-amyrin synthase

Leveraging a minimally engineered yeast strain that produces the precursor oleanolic acid at 1.35 g/L, we propose a strategy to align CYP450 expression with ethanol assimilation to enhance precursor supply and metabolic flux towards quillaic acid synthesis while reducing competition with growth-related processes. Additionally, we tuned CYP450 expression and selected efficient CPRs to further improve quillaic acid production.

## 2. Materials and methods

### 2.1 Plasmid and strain construction

Heterologous genes were codon-optimized and synthesized by either Twist Bioscience (South San Francisco, CA, USA) or Gene Universal (Newark, DE, USA) (Table S1). The target fragments were amplified by PCR. PCR products were analysed on 1% agarose gel, and the products with the correct size were purified with MinElute Gel Extraction Kit (QIAGEN, VIC, Australia) for further use. Genes involved in the synthesis pathway of quillaic acid were cloned into integrative vectors from EasyClone-MarkerFree Vector Set (ADDGENE, MA, USA) (Jessop-Fabre et al., 2016) through Gibson assembly using the NEBuilder® HiFi DNA Assembly Cloning Kit (New England Biolabs, Ipswich, MA, USA). Chemically competent *Escherichia coli* DH5α cells were prepared using the Mix & Go! *E. coli* Transformation Kit (Zymo Research, Irvine, CA, USA). Plasmids were extracted with Monarch® PCR & DNA Cleanup Kit (New England Biolabs, Ipswich, MA, USA). All yeast strains used in this study were derived from *S. cerevisiae* CEN.PK 102-5B (Euroscarf, Frankfurt am Main, Germany). Plasmids and linearized fragments were introduced into yeast by LiAc/ssDNA/PEG method (Daniel Gietz and Woods, 2002) (Table. S3). After incubating at 30 °C for 3 h, cells were plated on YPD agar with corresponding antibiotics. Positive transformants were identified by both colony PCR and Sanger sequencing of the PCR product.

### 2.2 Yeast shake-flask fermentation and intracellular triterpenoid extraction

A single colony from a fresh YPD agar plate was used to inoculate 5 mL of WM8+ mineral salt medium (Czarnotta et al., 2017) with 50 g/L glucose, incubated at 30°C and constant agitation of 250 rpm for 24 h. Cell growth was monitored by measuring the optical density at 600 nm (OD_600_) using an Avantor® Cell Density Meter CO8000. Cultures were diluted appropriately with fresh medium to ensure measurements remained within the linear range of detection (OD_600_ = 0.1 - 0.3). Prior to measurement, samples were vortexed to ensure homogeneity and measured against a blank of the corresponding fresh medium.

The pre-culture was used to inoculate 15 mL of WM8+ medium in a 250 mL shake-flask to an initial OD_600_ of 0.4. This main culture was incubated for 72 h at 30 °C and 250 rpm. A sample of 800 µL was taken and transferred into an Eppendorf tube prefilled with 250 µL 0.5 mm glass beads, 80 µL 5 M HCl and 800 µL methanol-chloroform (20:80). The sample underwent four cycles of intense agitation using a bead beater (Bead Mill Homogenizer, OMNI Inc., Kennesaw, GA, USA) at 4.5 m/s for 30 s each, with a 10 s pause between cycles to disrupt the cells. After centrifugation for 10 min at 17,000 x g, 500 µL of the lower chloroform phase were transferred into a high-performance liquid chromatography (HPLC) vial wit insert.

### 2.3 Metabolite analysis

Quantitative analysis of triterpenoids was performed using high-resolution liquid chromatography-mass spectrometry (LC-MS), with chromatographic separation conducted on an Exion UPLC system (Shimadzu, Kyoto, Japan; Pump A = 10% acetonitrile, 0.1% formic acid, and pump B = 90% acetonitrile, 0.1% formic acid). Samples (1 µl) were loaded onto a C8 column (2.1 x 150 mm, 100 A, 1.7 µm, Phenomenex, Torrance, CA, USA) at a flow rate of 400 µl/min in 3 % B for 0.5 min. Gradient consisted of 3% - 50% B over 12.5 min, followed by 50% - 97% B over 1 min, held at 97% B for 2.5 min and returned to 3% B over 0.5 min followed by 3 min re-equilibration. Eluted compounds were directly analysed on an X500B QTOF instrument (AB Sciex, Framingham, MA, USA) using a Sciex twin sprayer electrode. Source settings included CUR (curtain gas) = 25 psi, GS1 (ion Source Gas1) = 50 psi, GS2 (ion Source Gas2) = 50 psi, CAD (collision gas) = 7, temperature = 500°C and ion spray voltage = 5500 V. A TOF MS scan was performed across 100 - 2000 m/z for 0.125 sec followed by data-dependent acquisition. Workflow was set to “metabolite” with up to 20 molecules with +1 charge state and intensity greater than 200 counts/sec chosen for fragmentation using collision energy (CE) of 35 +/- 15 V. For MS1, DP = 50 V and CE = 5 V. MS2 acquisition was performed across 50-1800 m/z for 0.05 s. Samples were analysed in both positive and negative ion modes. For negative mode, the ion spray voltage was −4500 V, DP (declustering potential) = −50 V, and CE (collision energy) = −5 V (MS1) and CE-35 +/- 15 V with all other parameters the same.

Analytical standards of erythrodiol (97% purity; Sigma-Aldrich #09258), oleanolic acid (97% purity; Sigma-Aldrich #O5504), quillaic acid (98% purity; Cayman Chemical #31512), echinocystic acid (95% purity; Cayman Chemical #27476), hederagenin (97% purity; Sigma-Aldrich #H3916) and gypsogenin (Toronto Research Chemicals #G931808) were used to prepare the standard curves for quantification, ranging from 25 mg/L to 1 g/L (serial dilutions, dilution factor, df = 2).

Due to the lack of authentic standard of oleanolic aldehyde, the compound was purified from yeast triterpenoid extract using semi-preparative reverse-phase (RP)-HPLC. A binary gradient system consisting of acetonitrile and water with 0.1 % formic acid was employed on a C4 column (ACE 5 C4-300, 5 µm, 250 × 10 mm, Advanced Chromatography Technologies Ltd, Aberdeen, Scotland). The separation was performed with a gradient of 50% - 95 % acetonitrile (+0.1 % formic acid) over 47.5 min at a flow rate of 1 mL/min, with the column maintained at 45 °C. Compounds were detected at 214 nm using UV detection. The purified oleanolic aldehyde identified by NMR analysis (Table S4) was then utilized as a reference standard for quantification, consistent with previously mentioned standards. The specific titre was calculated by dividing the volumetric titre by the OD_600_ value.

Extracellular metabolites in the yeast cultures were analysed using HPLC on a Thermo Vanquish Duo system, equipped with both a diode array detector (DAD) set at 210 nm and a refractive index detector (RID) maintained at 40 °C. Separation was achieved using either an HiPlex H (H⁺ 8%, 300 × 7.7 mm, PN: PL1170-6830, Agilent Technologies, Santa Clara, CA, USA) or a Rezex RHM Monosaccharide column (H⁺ 8%, 300 × 7.8 mm, PN: 00H-0132-K0, Phenomenex, Torrance, CA, USA), with a SecurityGuard Carbo-H guard column (4 × 3 mm, PN: AJ0-4490, Phenomenex, Torrance, CA, USA) for sample protection. The column temperature was maintained at 65°C to optimize resolution and peak separation. Samples were injected at a volume of 25 µL, and separation was carried out using an isocratic mobile phase of 4 mM H₂SO₄ at a flow rate of 0.6 mL/min.

### 2.4 Real-time monitoring of yeast growth dynamics

The growth dynamics of yeast cultures were monitored using the Enzyscreen Growth Profiler 960 (Enzyscreen BV, Heemstede, The Netherlands). Yeast strains were cultivated in 24-deep-well plates (PN: CR1424d, Enzyscreen BV, Heemstede, The Netherlands), with each well containing 1.5 mL of culture. The plate was covered with a sandwich cover (PN: CR1224f, Enzyscreen BV, Heemstede, The Netherlands) to prevent contamination and minimize evaporation while allowing sufficient gas exchange. Prior to inoculation, overnight yeast cultures were adjusted to an OD_600_ of 0.05 and transferred to the wells in triplicate. The plates were placed into the Growth Profiler and incubated at 30 °C with orbital shaking at 250 rpm. Growth was recorded by measuring the backscatter signal every hour.

Raw image data were processed and analysed using the GP960 Viewer software. To convert backscatter signals (G value) into cell dry weight (CDW), a calibration curve specific to each strain in WM8+ medium between OD_600_ and G value was firstly established. It was carried out using a 16-point dilution series of each strain in the same medium, and a polynomial regression was fitted to the resulting data using the GP960 Viewer’s built-in tools. The generated equations were subsequently applied to convert backscatter measurements in experimental runs into estimated OD_600_ values. To relate OD₆₀₀ to CDW, standard curves were generated by measuring the dry weight of five different cell dilutions. Cell pellets were washed twice with 0.1 M NaCl and dried at 60 °C for 72 hours. To ensure complete drying, changes in dry weight were monitored for several additional hours until a constant weight was achieved. The growth curves for each strain were then generated based on the calculated CDW.

### 2.5 Proteomics analysis

To investigate protein expression dynamics during different yeast growth phases, we monitored the growth of the yeast strains in 24-well plates and identified five distinct growth phases (Fig. S1). Based on this profile, yeast samples were collected for proteomic analysis at two key time points: 13 h (exponential growth on glucose) and 38 h (post-diauxic phase on ethanol) for proteomics analysis.

Protein extraction was carried out using the S-Trap™ Micro Spin Column Digestion. Yeast cells were lysed by adding 25 µL of sodium dodecyl sulfate (SDS) lysis buffer to 25 µL of yeast cell suspension, incubated at 70 °C for 1 h, followed by overnight digestion at 37 °C. Samples were then sonicated for 10 minutes and centrifuged at 16,000 g for 5 min. A 10 µL aliquot was taken for protein quantification using a bicinchoninic acid (BCA) assay. For reduction and alkylation, samples were diluted to 50 µL, treated with 20 mM dithiothreitol at 70 °C for 1 h, cooled to room temperature, and incubated with 40 mM iodoacetamide in the dark for 15 minutes. Proteins were precipitated by acidification with 5 µL of 27.5 % phosphoric acid and bound to S-Trap columns using a binding buffer containing 90 % methanol and 100 mM Tris. Columns were washed three times before digestion with either trypsin alone (1 µg; 1:50 - 1:100 weight ratio) or sequential digestion with Lys-C (0.5 µg for 1 h) followed by trypsin (0.5 µg in two steps). Digestion was performed in 50 mM ammonium bicarbonate (ABC) buffer (pH 8.0) at 47°C for 1 h or overnight at 37°C. Peptides were eluted sequentially using increasing concentrations of acetonitrile in 0.1 % formic acid and pooled for drying and resuspension in 5 % acetonitrile with 0.1% formic acid for liquid chromatography-tandem mass spectrometry (LC-MS/MS) analysis.

Samples were then injected into a trap column (PepMap™ Neo Trap Cartridge, 22 mm × 300 µm, 5 µm C18, PN: 174500, Thermo Scientific, Waltham, MA, USA) at a flow rate of 10 µL/min, followed by separation on a resolving column (nanoEase™ column, 100 mm × 150 µm, 1.8 µm, 100 Å, Waters, Milford, MA, USA). Elution was achieved using a gradient of 8 % - 95 % ACN over 57 min. Mass spectrometry was performed using a Thermo Fisher Scientific UHPLC system coupled to an Exploris 480 mass spectrometer with a high-field asymmetric waveform ion mobility spectrometry (FAIMS) Pro interface. FAIMS compensation voltages were set at −45 V and −65 V. Full MS scans were acquired over an m/z range of 340 - 1110 at a resolution of 120,000. Tandem MS was performed in multiple isolation windows with a quadrupole isolation mode. Data analysis was conducted using Spectronaut software against a reference proteome that included both *Saccharomyces cerevisiae* CEN.PK proteome from the *Saccharomyces* Genome Database (SGD, https://www.yeastgenome.org/) and the proteins introduced into the strain. Differential protein abundance was assessed through pairwise comparisons across experimental groups defined by strain and growth phase. Comparisons with Q-values below 0.05 and valid ratio counts (≥8) were considered statistically significant. Valid ratio counts refer to the number of pairwise peptide ratios used to calculate the fold change between two conditions. For each protein, average group abundance within each group calculated from the normalized abundance values across biological triplicates, was used to derive log2 fold changes.

### 2.6 Fed-batch fermentation in bioreactors

A single colony of strain yQA14 was used to inoculate 5 mL of WM8+ mineral salt medium with 50 g/L glucose in 50 mL falcon tubes with (Czarnotta et al., 2017), and the cultures were incubated overnight at 30 °C and 250 rpm. Four 250 mL flasks with 15 mL fresh WM8+ medium were inoculated with the preculture to an OD_600_ of 0.4 and grown to an OD_600_ of 10 – 15 within 8 h. Subsequently, the cultures were transferred to a Multifors® II bench-top bioreactor (INFORS HT, Switzerland) containing 250 mL of WM8+ medium, adjusting the initial OD_600_ to 0.4. The pH was maintained at 6.0 using sodium hydroxide (NaOH) or phosphoric acid (H₃PO₄). Fermentations were conducted at 30 °C, with stirring and aeration controlled by a cascade system to maintain at dissolved oxygen (DO) level of 40 %. Ethanol feed, consisting of pure ethanol, was supplied at a rate of 2.1 mL/min to achieve a final concentration of 25 g/L. After an initial batch phase of 48 h, feeding was triggered whenever the DO level exceeded 60 %. After 72 h, additional WM8+ salts, trace elements and vitamins were added to the cell culture in the same amounts as those in the original medium to ensure sufficient availability for yeast growth. The whole process was controlled through eve® bioprocess platform software (INFORS HT, Switzerland).

## 3. Results and discussion

### 3.1 Constructing the quillaic acid pathway in yeast

The quillaic acid pathway was reconstructed in the *S. cerevisiae* chassis strain rOA2, previously engineered for oleanolic acid production (Guo et al., 2022). In this strain the MVA pathway was upregulated by expressing a truncated *HMG1* (Polakowski et al., 1998) and 2,3-oxidosqualene flux into ergosterol synthesis was reduced by replacing the native lanosterol synthase Erg7 with a destabilised variant (Guo et al., 2022) while oleanolic acid synthesis was enabled through the recombinant expression of an amyrin synthase from *Artemisia annua* (Kirby et al., 2008), C-28 oxidase CYP716A15 from *Vitis vinifera* (Fukushima et al., 2011) and a compatible reductase, MTR1, from *Medicago truncatula* (Young et al., 2011).

Three candidate genes each of C-16α and C-23 oxidases were individually expressed in rOA2 to evaluate their suitability for quillaic acid synthesis (Fig. 2a). C-16α oxidases sequentially oxidise the C16 of oleanolic acid (and related pentacyclic triterpenoids) into hederagenin and gypsogenin; candidates tested were CYP716-201209 (Q9) from *Quillaja saponaria*, CYP87D16 (M6) from *Maesa lanceolata* and CYP716A262 (P2) from *Psammosilene tunicoides*. Candidates of C-23 oxidases, which catalyse oxidation of the C-23 position of oleanolic acid (and analogous triterpenoid structures) to echinocystic acid, included CYP714-7 (Q7) from *Q. saponaria*, CYP72A68 (M8) from *M. truncatula* and CYP716A257 (P7) from *P. tunicoides* (Fukushima et al., 2013; Kulagina et al., 2021; Li et al., 2021; T. Liu et al., 2022; Moses et al., 2014). Catalytic efficiency was evaluated by individually expressing each gene under a constitutive promoter: pPGK1 for C-16α oxidases (Q9, P2, and M6) and pTEF1 for C-23 oxidases (Q7, P7, and M8) and quantifying triterpenoid production in shake flask experiments.

**Fig 2.**
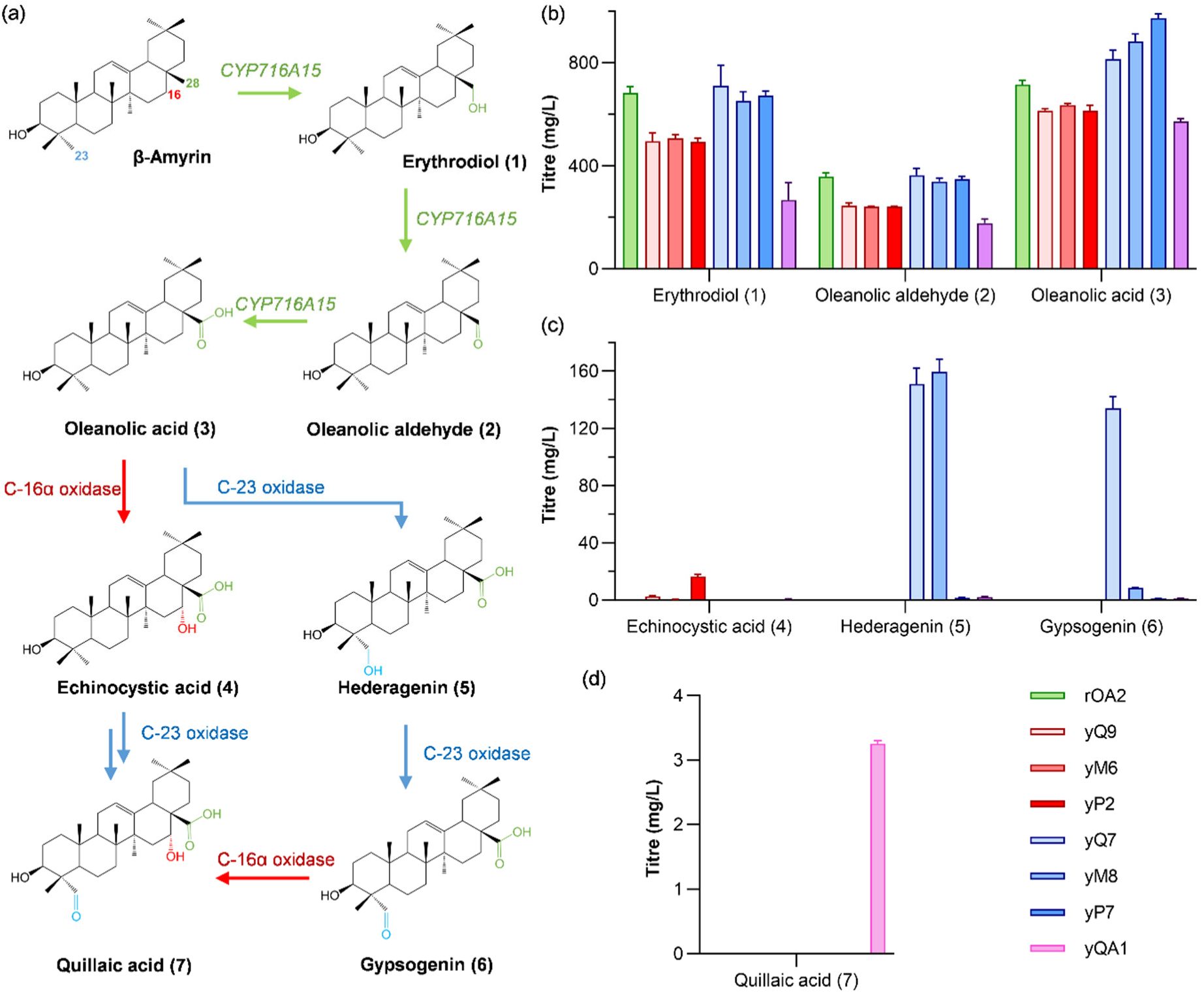
Identification of functional C-23 oxidases and C-16α oxidases for quillaic acid synthesis. (a) Quillaic acid biosynthesis from β-amyrin was implemented in strain rOA2 by expressing CYP450s candidates. (b-d) Triterpenoid quantification in yeasts expressing individual C-16α oxidases (yQ9, yM6, yP2) and C-23 oxidases (yQ7, yM8, yP7) and co-expressing CYP450s P2 and Q7 (yQA1). Data shown are the mean and standard deviation of single measurement from three shake-flask experiments. Relevant volumetric titres are provided in the main text.

The analysis of the products formed confirmed that only oleanolic acid (3) served as the substrate for these oxidases, while the oleanolic acid precursors, erythrodiol (1) and oleanolic aldehyde (2), which accumulated in the platform strain to a considerable extent, were not oxidised (Fig. 2a). Quantification revealed that P2 (C-16α oxidase) and Q7 (C-23 oxidase) produced the highest specific titres of echinocystic acid (4) and gypsogenin (6), respectively, with Q7 also producing hederagenin (5) (Fig. 2c). The corresponding genes of these enzymes were then co-expressed to establish the complete quillaic acid synthesis pathway in yeast. However, the resulting strain yQA1 produced only marginal titres of quillaic acid (7) (3.25 ± 0.05 mg/L) (Fig. 2d) despite an excess of the precursor oleanolic acid (3) (572 ± 10 mg/L) (Fig. 2b). Compared to the strains expressing the single CYP450s (yP2 and yQ7), yQA1 showed lower levels of echinocystic acid (4), hederagenin (5) and gypsogenin (6), suggesting that P2 and Q7 can further oxidize these intermediates to quillaic acid (7). This indicates that the low quillaic acid (7) production was caused by insufficient enzyme activity rather than substrate specificity of the CYP450s.

### 3.2 Carbon source dependent CYP450 expression for enhanced quillaic acid production

High residual titres of oleanolic acid (3) in yQA1 showed that P2 and Q7 activity limited quillaic acid (7) production and not the upstream oleanolic acid production. In yQA1, P2 and Q7 were controlled by the promoters pTEF1 and pPGK1, which are predominantly active during the initial glucose-driven fermentation stage (Sun et al., 2012). In order to increase the expression of *P2* and *Q7* in the post-diauxic growth phase, previously shown to favour triterpenoid production (Ebert et al., 2018), the promoters were exchanged with the galactose inducible pSkGAL2 and pSeGAL2 promoters. In addition, the Gal80 repressor was deleted in this strain (yQA2) so that these GAL promoters are auto-induced after glucose depletion (Peng et al., 2018; Torchia et al., 1984). Quillaic acid production was considerably improved in the resulting strain yQA2, with an increase of the quillaic acid titre by more than 80-fold to 279 ± 1 mg/L and an increase of all triterpenoids produced from 1104 ± 79 mg/L to 1452 ± 44 mg/L (Fig. 3a). No growth inhibition was observed in yQA2, indicating that the enhanced CYP450 expression did not impose a major burden (Fig. S1), potentially due to shift of expression to the ethanol phase.

**Fig 3.**
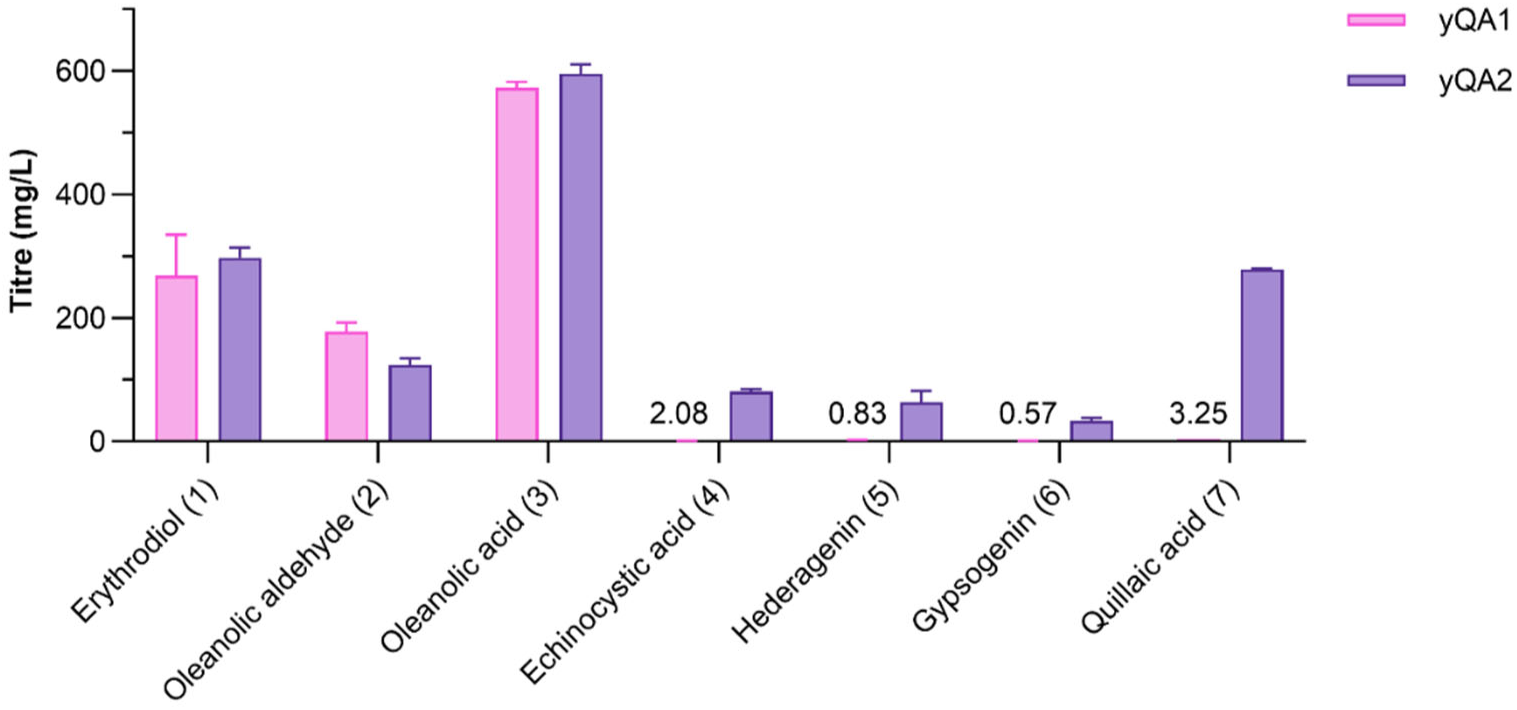
Metabolic analysis of yeasts expressing *CYP450*s under different promoters. Triterpenoid production in *S. cerevisiae* strains expressing P2 and Q7 using constitutive pPGK1 and pTEF1 promoters (yQA1) and (auto-)inducible promoters pSkGAL2 and pSeGAL2 (yQA2), respectively. Products were quantified after 72 h of flask cultivation. Data shown are the mean and standard deviation of single measurement from triplicate shake-flask experiments.

LC-MS analysis of the yQA2 yeast extract revealed a new compound with an m/z value of 487.34 [M– H]⁻ (Fig. S2), which agrees closely with the calculated mass (487.3429) for 16α-hydroxy-hederagenin. The compound was isolated and further characterized by NMR spectroscopy, confirming its identity as 16α-hydroxy-hederagenin (Table. S6). This intermediate can be further be oxidized at C-23 by Q7 to yield quillaic acid.

### 3.3 Proteomic changes to recombinant triterpenoid biosynthesis

We performed global proteomics analysis for yQA1 and yQA2 during the glucose and ethanol phase to assess differences in their metabolic responses (Fig. 4). Lanosterol synthase Erg7 was not detected in the proteomics data due to the low abundance of the destabilised mutant (Guo et al., 2022). As expected, P2, Q7 were significantly upregulated in yQA2 at ethanol phase compared to glucose phase, as expected due the auto-induction of GAL promoters, and CYP716A15 was also upregulated in yQA2 at ethanol phase despite being under control of the constitutive pTEF1 promoter in both strains.

**Fig 4.**
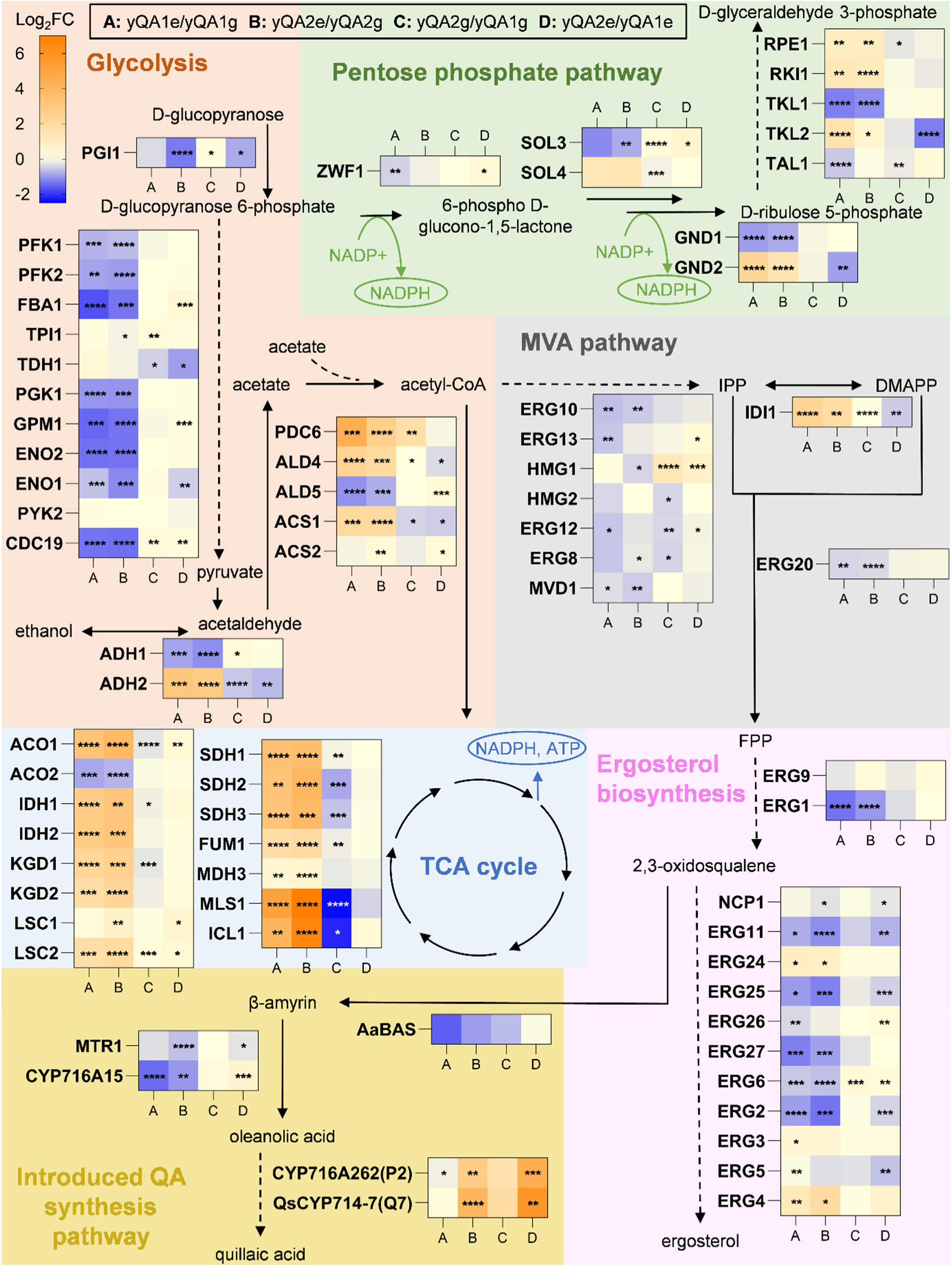
Heatmap depicting the protein expression involved in quillaic acid synthesis and related pathways in yQA1 and yQA2 strains. The heatmap shows log2 fold changes comparing various strains and growth phases, labelled as A (yQA1e/yQA1g), B (yQA2e/yQA2g), C (yQA2g/yQA1g), and D (yQA2e/yQA1e). yQA1g and yQA2g represent the exponential growth phase utilizing glucose as the carbon source, while yQA1e and yQA2e correspond to the post-diauxic growth phase utilizing ethanol. Proteomics analysis was conducted with three replicates. Significant differences are indicated by asterisks (*P < 0.05, **P < 0.01, ***P < 0.001, ****P < 0.0001).

During ethanol phase, several ergosterol-synthesis enzymes were downregulated, with stronger repression in yQA2 (Fig. 4). Meanwhile, ER membrane-biogenesis proteins (Scs2p, Scs3p, Cho1p and Dpm1p), the key phospholipid biosynthesis enzyme Ino1p, the chaperone Kar2p, and regulator Ire2p to unfolded protein response (UPR) were all elevated, indicating activation of the ER stress response. Consistent with our observations, ER expansion in response to the overexpression of CYP450 and CPR previously been reported previously (Zimmer et al., 2000). As ER membrane area appears to limit CYP/CPR expression, deliberately expanding the ER, an approach previously shown to enhance triterpenoid biosynthesis (Arendt et al., 2017), could further improve production efficiency.

In addition, pantothenate kinase Cab1p, which catalyses the first and rate-limiting step in Coenzyme A (CoA) biosynthesis (Olzhausen et al., 2009), was upregulated in yQA2 during the ethanol phase. Enhanced CoA availability could support higher flux through the MVA pathway, consistent with the previous report demonstrating that enhanced CoA synthesis can improve acetyl-CoA-derived product formation (Schadeweg and Boles, 2016). Meanwhile, enzymes involved in competing long-chain fatty acid biosynthesis (Fas2p, Faa1p, Ifa38p, Scs7p) were downregulated in yQA2, suggesting endogenous re-allocation of precursors and cofactors such as acetyl-CoA and NADPH toward triterpenoid biosynthesis(Jin et al., 2019; Li and Zhang, 2004).

### 3.4 Optimizing CYP450 and CPR ratios for productivity

To further improve quillaic acid production, additional gene copies of *P2* and *Q7* were introduced and expressed under the control of GAL2 promoters, resulting in strain yQA3 (2 copies) and yQA4 (3 copies). While yQA4 showed a 1.7-fold increase in specific quillaic acid titre (g/L per OD_600_), its growth was impaired and the final OD_600_ after 72 h fermentation dropped to 65 (72% of final OD of yQA2) and no improvement in volumetric titre of quillaic acid was observed (Fig. 5a).

**Fig 5.**
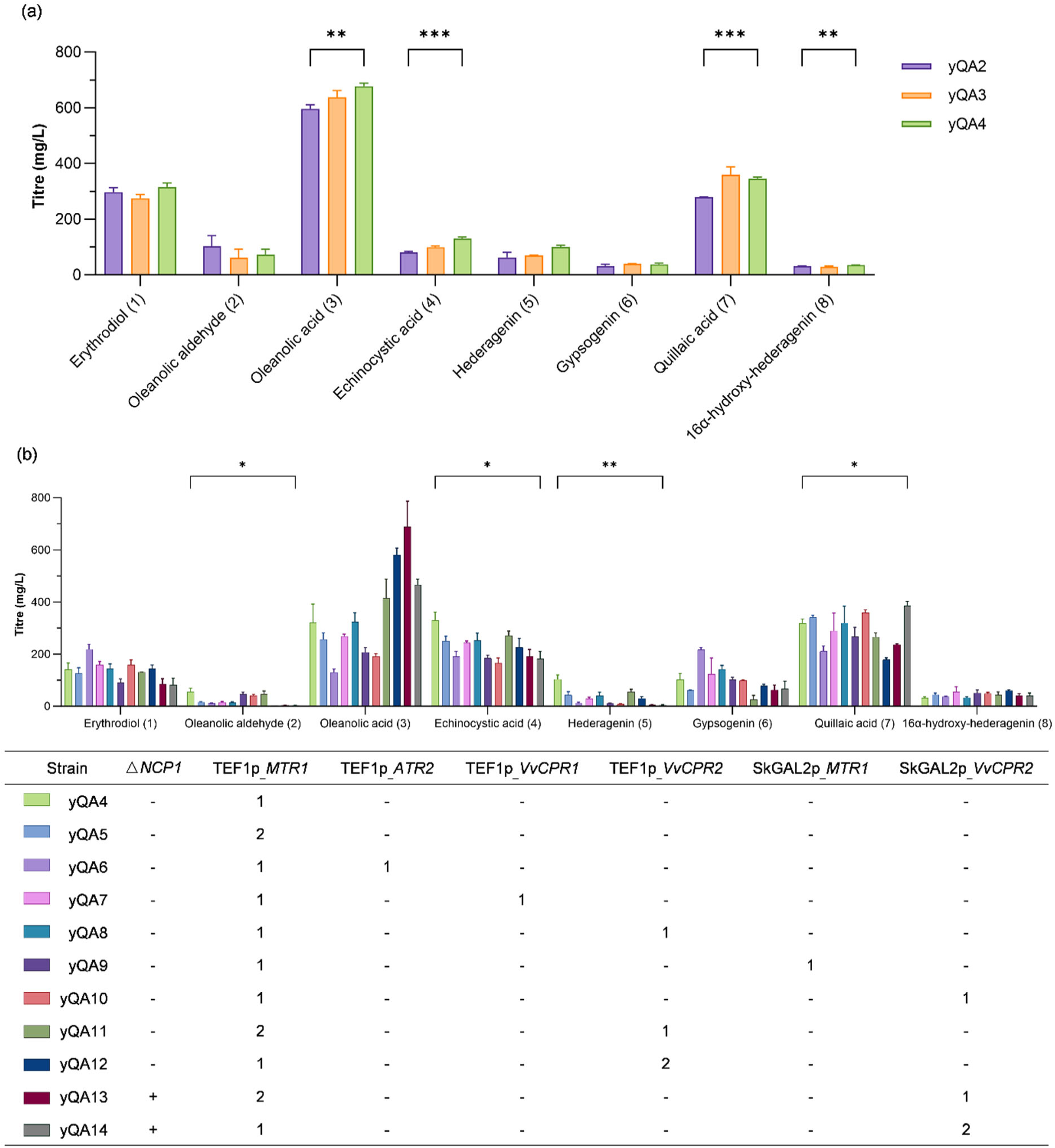
Triterpenoid production in yeast with varied expression ratio of CYP450s and CPRs. (a) Titres of target triterpenoids in yeasts expressing 1 - 3 copies of *P2* and *Q7* by GAL2 promoters. (b) Screening of CPRs and fine-tuning the expression level. Four CPR candidates were expressed with different promoters in yQA4 strain. Data shown are the mean and standard deviation of measurements from triplicate shake-flask experiments.

To identify CPRs that optimally transfer electrons to the CYP450s involved in quillaic acid synthesis, four candidates were expressed individually in yQA4 under pTEF1 control: MTR1 from *M. truncatula* and ATR2 from *Arabidopsis thaliana*, commonly used to support plant derived P450 activity in yeast (Fig. 5b), as well as the CYP716A15 cognate VvCPRs, to test whether CYP450s function more efficiently with reductases from the same origin, as observed for some CYP450s (Jennewein et al., 2005). Additionally, to compare reductases from different functional classes, two class I (MTR1 and VvCPR1) and two class II (ATR2 and VvCPR2) CPRs were included.

Compared to yQA4, with a single MTR1 copy, adding an extra copy of either MTR1 or VvCPR2 increased quillaic acid titre. However, further increasing the copy number of these genes led to reduced quillaic acid titre and increased oleanolic acid (3) accumulation, particularly in strain yQA12, which expressed two copies of *MTR1* and two copies of *VvCPR2*. This suggests that the ratio of CYP450 to CPR is a key determinant of product distribution. In fact, the growth of the strain expressing three copies of *MTR1*, was completely inhibited in mineral salt medium.

To fine-tune CYP450s:CPRs stoichiometry, we placed MTR1 or VvCPR2 under GAL promoter control. MTR1 performed best when constitutively expressed from pTEF1 (yQA5), resulting in higher quillaic acid titre than under pSkGAL2 (yQA9). In contrast, *VvCPR2* gave the highest quillaic acid triter when expressed under pSkGAL2 control (yQA10), with even more significant improvement when considering the quillaic acid proportion among total triterpenoids (Fig. S5b). A reduced fraction of hederagenin (5) and gypsogenin (6) in yQA10 than in yQA8 suggests that the pSkGAL2-based expression improved the ratio of VvCPR2 and P2, facilitating electron transfer and hence conversion of intermediates into quillaic acid. This also explains why more oleanolic acid (3) was produced when expressing two copies of *VvCPR2*, as P2 is a multifunctional enzyme capable of oxidizing the C-28 and C-16 carbon atoms.

We further deleted the yeast’s endogenous CPR (encoded by *NCP1*) to determine its impact on heterologous CYP450 activity. To maintain a consistent CPR copy number after *NCP1* deletion, an additional copy of either *MTR1* or *VvCPR2* was expressed under the promoter that previously supported higher quillaic acid production, generating strains yQA13 and yQA14. yQA14, in which *NCP1* was deleted and two copies of *VvCPR2* were expressed under pSkGAL2 control along with the existing *MTR1*, achieved the highest overall performance, producing a maximum quillaic acid titre of 385 ± 17 mg/L (Fig. 5b), along with the highest specific titre (Fig. S5a) and the second greatest proportion of quillaic acid (Fig. S5b) among total triterpenoids, indicating that VvCPR2 is the most effective CPR among the candidates for quillaic acid synthesis in the engineered yeast. However, Ncp1p (in yQA10) proved more effective than MTR1 (in yQA13) for quillaic acid production, while yQA13 produced more oleanolic acid (3). Considering the high accumulation of oleanolic acid and echinocystic acid (4), the follow up work should focus on exploring either more selective C28 oxidases and C16α oxidases with broader substrate flexibility and higher conversion efficiency.

### 3.5 Fed-Batch fermentation for maximizing titre

To scale up the quillaic acid production under more controlled environmental conditions and with increased carbon source and oxygen availability, yQA4, which performed best in shake flask experiments, was cultivated in triplicate fed-batch bioreactor experiments (BR1, BR2, BR3), which demonstrated the best overall performance in flask-based quillaic acid production. Carbon source was switched to ethanol after an initial glucose batch phase.

Ethanol feeding during the fed-batch fermentation was initiated when DO levels exceeded 60% (Fig. S6a), indicating carbon source depletion, with each ethanol pulse replenishing ethanol to a concentration of approximately 25 g/L. Although the same preculture and identical fermentation conditions were used, differences in cell growth and triterpenoid production were observed, likely due to unavoidable technical variabilities between individual bioreactors. BR2 consistently showed the highest quillaic acid production throughout the fed-batch fermentation (Fig. S6b) and was therefore selected for further detailed analysis. Among the multiple ethanol additions in BR2, only the initial ethanol pulse notably increased the quillaic acid titre to a maximum of 471 ± 20 mg/L (Fig. 6a) recorded after around 81 h. At this point of the fed-batch fermentation, quillaic acid was the dominant product, accounting for 65 % of the quantified triterpenoids compared to 32.6 % in flask cultures (Fig. 6b). Enhanced oxygen supply and potentially improved NADPH provision promoted the oxidation of oleanolic acid (3) to quillaic acid. This indicates that fermentation process optimization is as critical as strain engineering for maximizing biosynthetic efficiency.

**Fig 6.**
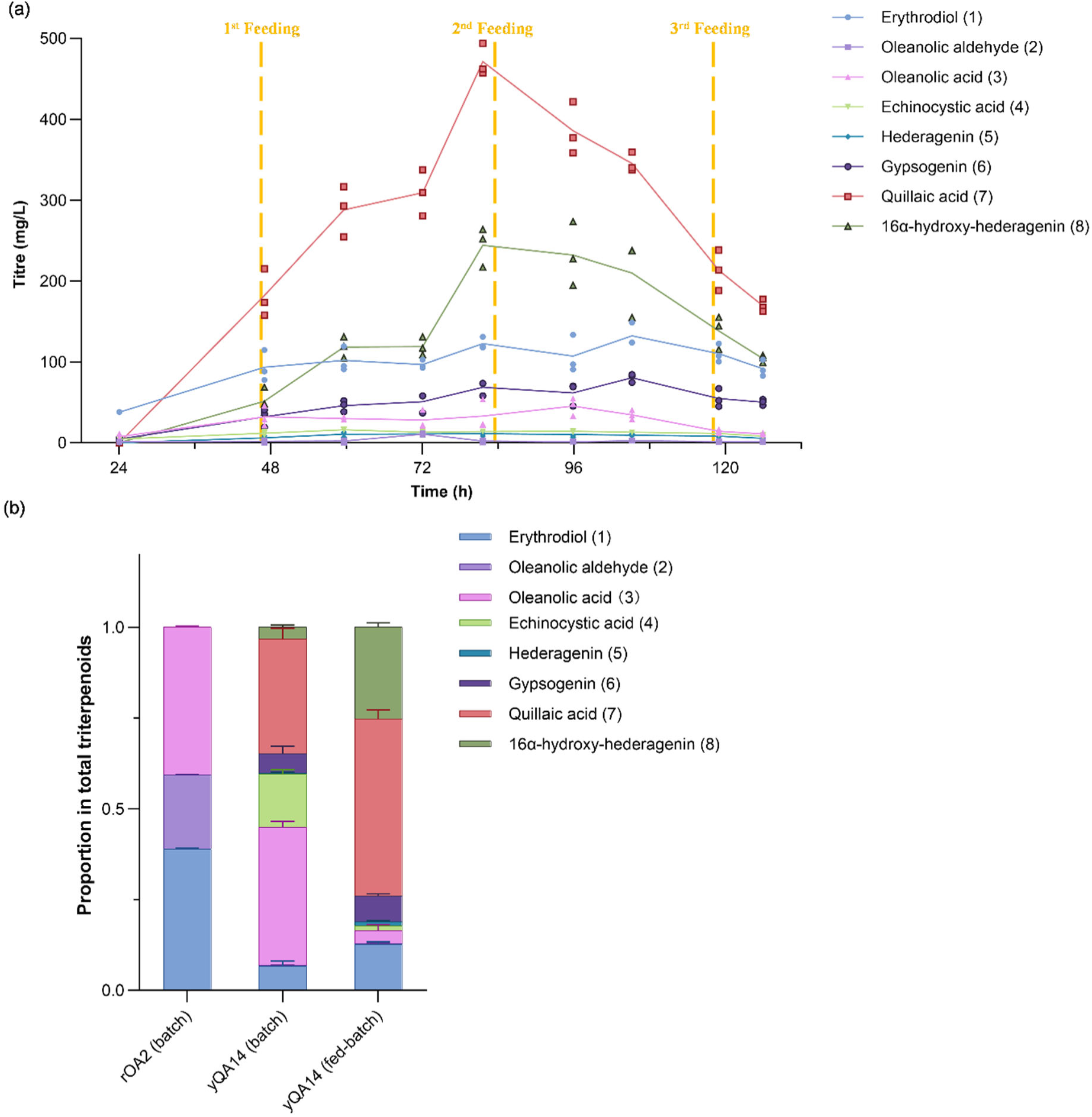
Scale-up of triterpenoid production through fed-batch fermentation in bioreactors. (a) Triterpenoid production in strain yQA14 during fed-batch fermentation. The lines represent the mean values, and the individual symbols correspond to the single measurements from triplicate bioreactor experiments. (b) Comparison of the triterpenoid distribution extracted from the chassis strain (rOA2) and yQA14 after 72 h of flask fermentation and 81.5 h of fed-batch fermentation in a bioreactor, respectively.

In the further course of the fermentation, quillaic acid unexpectedly decreased to less than 50 % of its maximum titre within approximately 48 h. The 16α-hydroxy hederagenin (8) concentration followed a similar trend to quillaic acid (Fig. S6d). Similar decreasing trend was also observed in the other two bioreactors (Fig. S6b, c). Such a behaviour has not been observed in our previous work where upscaling from shake flask to bioreactor experiments resulted in a constant increase over the fermentation and a five-fold overall improvement of triterpenoid production (Guo et al., 2022).

Two additional new peaks (NP1 and NP2) were detected in LC-MS analysis. NP1 accumulated during the early stages of fermentation, showed a slight decline over time, and abruptly disappeared around 120 h. At this point, NP2 emerged and began to increase after the third ethanol feeding (Fig. S6b), indicating that its synthesis may be associated with the decline in quillaic acid production. Future work will focus on understanding these metabolite dynamics to improve quillaic production.

Although increases in stirrer speed and O_2_ flow were observed after each feeding to sustain the DO setpoint, the O_2_ flow increment after the second feeding (136.32 L) was smaller compared to the first (143.34 L). This pattern suggests that, despite rising cell density, the oxygen consumption per cell biomass decreased. Concurrently, organic acid analysis of the cultural medium revealed that acetate began accumulating during the later stages of fermentation, indicating that ethanol was not effectively directed toward triterpenoid synthesis but instead diverted into acetate, reaching a concentration of 14.17 mM by 118 h (Fig. S7a) (Paczia et al., 2012). The combined observations of reduced specific oxygen uptake and increased acetate accumulation can be a sign of limited TCA cycle activity or inhibition of respiration. This could reflect a broader metabolic stress, potentially linked to lipotoxicity caused by triterpenoid accumulation in (sub-)cellular membranes.

## 4. Conclusion

CYP450s play a crucial role in triterpenoid synthesis (Dinday & Ghosh, 2023). To enhance their functional expression in yeast, researchers have employed various strategies, including modifying CYP450 protein structures (Romsuk et al., 2022) and optimising other elements within the reaction environment (Cha et al., 2022; Chen et al., 2022; Iizaka et al., 2021). In our study, we demonstrate that CYP450 activity is influenced not only by expression level but also by the timing of expression during yeast growth. Selecting appropriate CYP450/CPR combinations and adjusting expression ratios improved production and altered product distribution, however, gains were limited by the challenge of identifying CPRs that are broadly compatible with multiple CYP450s. High-throughput combinatorial screening could address this limitation by more efficiently uncovering optimal pairings.

Fed-batch fermentation in bioreactors revealed optimizing fermentation conditions is just as critical as genetic engineering for improving the production of target compounds in microbial synthesis systems. Because quillaic acid became unstable and converted to other metabolites in the late in fed-batch phase, enhancing product stability will be essential to achieve higher titres and consistent yields.

## Acknowledgements

We gratefully acknowledge the Queensland Node of Metabolomics and Proteomics Australia (Q-MAP) for their support in acquiring metabolomics and proteomics data, with special appreciation to Dr Gabi Netzel for her assistance with metabolomics data acquisition. Q-MAP is supported by Bioplatforms Australia, an NCRIS-funded initiative. We also thank Integrated Design Environment for Advanced biomanufacturing (IDEA Bio) for providing access to their benchtop bioreactor systems, especially Dr. Axayacatl Gonzalez and Ms. Yu Sun for their invaluable technical support, training, and guidance throughout the bioreactor operation. This work was supported by the Australian Government Department of Education through the Trailblazer Universities Program “Food and Beverage Accelerator (FaBA)”, and the China Scholarship Council (CSC) (No. 202104910075).

## References

Arendt, P., Miettinen, K., Pollier, J., De Rycke, R., Callewaert, N., Goossens, A., 2017. An endoplasmic reticulum-engineered yeast platform for overproduction of triterpenoids. Metab Eng, 40, 165–175. 10.1016/j.ymben.2017.02.007

Brown, S., Clastre, M., Courdavault, V., O’Connor, S. E., 2015. De novo production of the plant-derived alkaloid strictosidine in yeast. Proc. Natl. Acad. Sci., 112(11), 3205–3210. 10.1073/pnas.1423555112

Cha, Y., Li, W., Wu, T., You, X., Chen, H., Zhu, C., Zhuo, M., Chen, B., Li, S., 2022. Probing the synergistic ratio of p450/cpr to improve (+)-nootkatone production in *Saccharomyces cerevisiae*. J. Agric. Food Chem., 70(3), 815–825. 10.1021/acs.jafc.1c07035

Czarnotta, E., Dianat, M., Korf, M., Granica, F., Merz, J., Maury, J., Baallal Jacobsen, S. A., Förster, J., Ebert, B. E., Blank, L. M., 2017. Fermentation and purification strategies for the production of betulinic acid and its lupane-type precursors in *Saccharomyces cerevisiae*. Biotechnol. Bioeng., 114(11), 2528–2538. 10.1002/bit.26377

Daniel Gietz, R., Woods, R. A. (2002). Transformation of yeast by lithium acetate/single-stranded carrier DNA/polyethylene glycol method, in: Guthrie, G., Fink, G. R. (Eds), Methods in Enzymology. Academic Press, the U. S., pp. 87–96. 10.1016/S0076-6879(02)50957-5

Dinday, S., Ghosh, S., 2023. Recent advances in triterpenoid pathway elucidation and engineering. Biotechnol. Adv., 68, 108214–108214. 10.1016/j.biotechadv.2023.108214

Ebert, B. E., Czarnotta, E., Blank, L. M., 2018. Physiologic and metabolic characterization of *Saccharomyces cerevisiae* reveals limitations in the synthesis of the triterpene squalene. FEMS Yeast Res., 18(8). 10.1093/femsyr/foy077

Emmerstorfer, A., Wimmer-Teubenbacher, M., Wriessnegger, T., Leitner, E., Muller, M., Kaluzna, I., Schurmann, M., Mink, D., Zellnig, G., Schwab, H., Pichler, H., 2015. Over-expression of *ICE2* stabilizes cytochrome P450 reductase in *Saccharomyces cerevisiae* and *pichia pastoris*. Biotechnol. J., 10(4), 623–635. 10.1002/biot.201400780

Fukushima, E. O., Seki, H., Sawai, S., Suzuki, M., Ohyama, K., Saito, K., Muranaka, T., 2013. Combinatorial biosynthesis of legume natural and rare triterpenoids in engineered yeast. Plant Cell Physiol., 54(5), 740–749. 10.1093/pcp/pct015

Garçon, N., Van Mechelen, M., 2011. Recent clinical experience with vaccines using MPL- and QS-21-containing adjuvant systems. Expert Rev. Vaccines, 10(4), 471–486. 10.1586/erv.11.29

Guo, H., Jacobsen, S. A. B., Walter, K., Lewandowski, A., Czarnotta, E., Knuf, C., Polakowski, T., Maury, J., Lang, C., Förster, J., Blank, L. M., Ebert, B. E., 2022. Triterpenoid production with a minimally engineered *Saccharomyces cerevisiae* chassis. bioRxiv., 2022.2007.2011.499565. 10.1101/2022.07.11.499565

Guo, H., Wang, H., Huo, Y.-X., 2020. Engineering critical enzymes and pathways for improved triterpenoid biosynthesis in yeast. ACS Synth. Biol., 9(9), 2214–2227. 10.1021/acssynbio.0c00124

Istiandari, P., Yasumoto, S., Srisawat, P., Tamura, K., Chikugo, A., Suzuki, H., Seki, H., Fukushima, E. O., Muranaka, T., 2021. Comparative analysis of NADPH-cytochrome P450 reductases from legumes for heterologous production of triterpenoids in transgenic *Saccharomyces cerevisiae*. Front. Plant Sci., 12. 10.3389/fpls.2021.762546

Jennewein, S., Park, H., DeJong, J. M., Long, R. M., Bollon, A. P., Croteau, R. B., 2005. Coexpression in yeast of taxus cytochrome P450 reductase with cytochrome P450 oxygenases involved in taxol biosynthesis. Biotechnol. Bioeng., 89(5), 588–598. 10.1002/bit.20390

Jessop-Fabre, M. M., Jakociunas, T., Stovicek, V., Dai, Z., Jensen, M. K., Keasling, J. D., Borodina, I., 2016. Easyclone-Markerfree: A vector toolkit for marker-less integration of genes into *Saccharomyces cerevisiae* via CRISPR-Cas9. Biotechnol. J., 11(8), 1110–1117. 10.1002/biot.201600147

Jiang, L., Huang, L., Cai, J., Xu, Z., Lian, J., 2021. Functional expression of eukaryotic cytochrome P450s in yeast. Biotechnol. Bioeng., 118(3), 1050–1065. 10.1002/bit.27630

Jin, C.-C., Zhang, J.-L., Song, H., Cao, Y.-X., 2019. Boosting the biosynthesis of betulinic acid and related triterpenoids in *Yarrowia Lipolytica* via multimodular metabolic engineering. Microb. Cell Fact., 18(1), 77. 10.1186/s12934-019-1127-8

Jobe, O., Kim, J., Pinto, D. O., Villar, Z., Hewitt, T., Duncan, E. H., Anderson, A., Gohain, N., Gong, H., Tucker, C., Alving, C. R., Matyas, G. R., Bergmann-Leitner, E., Rao, M., 2022. Army liposome formulation containing QS-21 render human monocyte-derived macrophages less permissive to HIV-1 infection by upregulating APOBEC3A. Sci. Rep., 12(1), 7570–7570. 10.1038/s41598-022-11230-8

Kim, J. E., Jang, I. S., Sung, B. H., Kim, S. C., Lee, J. Y., 2018. Rerouting of NADPH synthetic pathways for increased protopanaxadiol production in *Saccharomyces cerevisiae*. Sci. Rep., 8(1), 15820. 10.1038/s41598-018-34210-3

Kulagina, N., Guirimand, G., Melin, C., Lemos-Cruz, P., Carqueijeiro, I., De Craene, J.-O., Oudin, A., Heredia, V., Koudounas, K., Unlubayir, M., Lanoue, A., Imbault, N., St-Pierre, B., Papon, N., Clastre, M., Giglioli-Guivarc’h, N., Marc, J., Besseau, S., Courdavault, V., 2021. Enhanced bioproduction of anticancer precursor vindoline by yeast cell factories. Microb. Biotechnol., 14(6), 2693–2699. 10.1111/1751-7915.13898

Li, J., Zhang, Y., 2014. Increase of betulinic acid production in *Saccharomyces cerevisiae* by balancing fatty acids and betulinic acid forming pathways. Appl. Microbiol. Biotechnol., 98(7), 3081–3089. 10.1007/s00253-013-5461-1

Li, W. X., Ma, X. H., Li, G. D., Zhang, A. L., Wang, D., Fan, F. Y., Ma, X. L., Zhang, X. L., Dai, Z. B., Qian, Z. G., 2021. De novo biosynthesis of the oleanane-type triterpenoids of tunicosaponins in yeast. ACS Synth. Biol., 10(8), 1874–1881. 10.1021/acssynbio.1c00065

Liu, J., Wang, X., Dai, G., Zhang, Y., Bian, X., 2022. Microbial chassis engineering drives heterologous production of complex secondary metabolites. Biotech. Adv., 59, 107966–107966. 10.1016/j.biotechadv.2022.107966

Liu, T., Gou, Y., Zhang, B., Gao, R., Dong, C., Qi, M., Jiang, L., Ding, X., Li, C., Lian, J., 2022. Construction of ajmalicine and sanguinarine de novo biosynthetic pathways using stable integration sites in yeast. Biotechnol. Bioeng., 119(5), 1314–1326. 10.1002/bit.28040

Liu, Y., Zhao, X., Gan, F., Chen, X., Deng, K., Crowe, S. A., Hudson, G. A., Belcher, M. S., Schmidt, M., Astolfi, M. C. T., Kosina, S. M., Pang, B., Shao, M., Yin, J., Sirirungruang, S., Iavarone, A. T., Reed, J., Martin, L. B. B., El-Demerdash, A., Kikuchi, S., Misra, R. C., Liang, X., Cronce, M. J., Chen, X., Zhan, C., Kakumanu, R., Baidoo, E. E. K., Chen, Y., Petzold, C. J., Northen, T. R., Osbourn, A., Scheller, H., Keasling, J. D., 2024. Complete biosynthesis of QS-21 in engineered yeast. Nature, 629(8013), 937–944. 10.1038/s41586-024-07345-9

Modi, A. R., Dawson, J. H. (2015). Oxidizing intermediates in p450 catalysis: A case for multiple oxidants, in E. G. Hrycay, S. M. Bandiera (Eds.), Monooxygenase, peroxidase and peroxygenase properties and mechanisms of cytochrome P450. Springer International Publishing, pp. 63–81. 10.1007/978-3-319-16009-2_2

Moses, T., Pollier, J., Faizal, A., Apers, S., Pieters, L., Thevelein, J. M., Geelen, D., Goossens, A., 2014. Unravelling the triterpenoid saponin biosynthesis of the african shrub *Maesa Lanceolata*. Mol. Plant. 10.1093/mp/ssu110

Nowrouzi, B., Li, R. A., Walls, L. E., d’Espaux, L., Malcı, K., Liang, L., Jonguitud-Borrego, N., Lerma-Escalera, A. I., Morones-Ramirez, J. R., Keasling, J. D., Rios-Solis, L., 2020. Enhanced production of taxadiene in *Saccharomyces cerevisiae*. Microb. Cell Fact., 19(1), 200–200. 10.1186/s12934-020-01458-2

Olzhausen, J., Schübbe, S., Schüller, H. J., 2009. Genetic analysis of coenzyme a biosynthesis in the yeast *Saccharomyces cerevisiae*: Identification of a conditional mutation in the pantothenate kinase gene CAB1. Curr. Genet., 55(2), 163–173. 10.1007/s00294-009-0234-1

Özverel, C. S., Uyanikgil, Y., Karaboz, İ., Nalbantsoy, A., 2020. Investigation of the combination of anti-PD-L1 mAb with HER2/neu-loaded dendritic cells and QS-21 saponin adjuvant: effect against HER2 positive breast cancer in mice. Immunopharmacol. Immunotoxicol., 42(4), 346–357. 10.1080/08923973.2020.1775644

Paczia, N., Nilgen, A., Lehmann, T., Gätgens, J., Wiechert, W., Noack, S., 2012. Extensive exometabolome analysis reveals extended overflow metabolism in various microorganisms. Microb. Cell Fact., 11(1), 122. 10.1186/1475-2859-11-122

Papagiannidis, D., Bircham, P. W., Luchtenborg, C., Pajonk, O., Ruffini, G., Brugger, B., Schuck, S., 2021. ICE2 promotes er membrane biogenesis in yeast by inhibiting the conserved lipin phosphatase complex. EMBO J., 40(22), e107958. 10.15252/embj.2021107958

Peng, B., Wood, R. J., Nielsen, L. K., Vickers, C. E., 2018. An expanded heterologous *GAL* promoter collection for diauxie-inducible expression in *Saccharomyces cerevisiae*. ACS Synth. Biol., 7(2), 748–751. 10.1021/acssynbio.7b00355

Qu, Y., Easson, M. L. A. E., Froese, J., Simionescu, R., Hudlicky, T., Luca, V. D., 2015. Completion of the seven-step pathway from tabersonine to the anticancer drug precursor vindoline and its assembly in yeast. Proc. Natl. Acad. Sci., 112(19), 6224–6229. 10.1073/pnas.1501821112

Quintanilha, J. C. F., de Sousa, V. M., Visacri, M. B., Amaral, L. S., Santos, R. M. M., Zambrano, T., Salazar, L. A., Moriel, P., 2017. Involvement of cytochrome P450 in cisplatin treatment: Implications for toxicity. Cancer Chemother Pharmacol., 80(2), 223–233. 10.1007/s00280-017-3358-x

Ragupathi, G., Gardner, J. R., Livingston, P. O., Gin, D. Y., 2011. Natural and synthetic saponin adjuvant QS-21 for vaccines against cancer. Expert Rev. Vaccines, 10(4), 463–470. 10.1586/erv.11.18

Schadeweg, V., Boles, E., 2016. n-Butanol production in Saccharomyces cerevisiae is limited by the availability of coenzyme A and cytosolic acetyl-CoA. Biotechnol. Biofuels, 9, 44. 10.1186/s13068-016-0456-7

Srinivasan, P., Smolke, C. D., 2020. Biosynthesis of medicinal tropane alkaloids in yeast. Nature, 585(7826), 614–619. 10.1038/s41586-020-2650-9

Sun, M., Xin, Q., Hou, K., Qiu, J., Wang, L., Chao, E., Su, X., Zhang, X., Chen, S., Wang, C., 2023. Production of 11-oxo-beta-amyrin in *Saccharomyces cerevisiae* at high efficiency by fine-tuning the expression ratio of CYP450:CPR. J Agric Food Chem, 71(8), 3766–3776. 10.1021/acs.jafc.2c08261

Sun, J., Shao, Z., Zhao, H., Nair, N., Wen, F., Xu, J.-H., Zhao, H., 2012. Cloning and characterization of a panel of constitutive promoters for applications in pathway engineering in *Saccharomyces cerevisiae*. Biotechnol. Bioeng., 109(8), 2082–2092. 10.1002/bit.24481

Sun, Z.-J., Lian, J.-Z., Zhu, L., Jiang, Y.-Q., Li, G.-S., Xue, H.-L., Wu, M.-B., Yang, L.-R., Lin, J.-P., 2021. Combined biosynthetic pathway engineering and storage pool expansion for high-level production of ergosterol in industrial *Saccharomyces cerevisiae*. F Front. Bioeng. Biotechnol., 9. 10.3389/fbioe.2021.681666

Wang, P., Škalamera, Đ., Sui, X., Zhang, P., Michalek, S. M., 2019. Synthesis and evaluation of QS-7-based vaccine adjuvants. ACS Infect. Dis., 5(6), 974–981. 10.1021/acsinfecdis.9b00039

Wang, Y. C., Chen, C. R., Chen, C. Y., Liang, P. H., 2024. Synthesis of quillaic acid through sustainable C-H bond activations. J. Org. Chem., 89(8), 5491–5497. 10.1021/acs.joc.3c02958

Yang, J., Liu, Y., Zhong, D., Xu, L., Gao, H., Keasling, J. D., Luo, X., Chou, H. H., 2023. Combinatorial optimization and spatial remodeling of CYPs to control product profile. Metab. Eng., 80, 119–129. 10.1016/j.ymben.2023.09.004

Zhou, A., Zhou, K., Li, Y., 2021. Rational design strategies for functional reconstitution of plant cytochrome P450s in microbial systems. Curr. Opin. Plant Biol., 60, 102005–102005. 10.1016/j.pbi.2021.102005

Zhu, Y., Li, J., Peng, L., Meng, L., Diao, M., Jiang, S., Li, J., Xie, N., 2022. High-yield production of protopanaxadiol from sugarcane molasses by metabolically engineered *Saccharomyces cerevisiae*. Microb. Cell Fact., 21(1), 230. 10.1186/s12934-022-01949-4

Zimmer, T., Ogura, A., Takewaka, T., Zimmer, R.-M., Ohta, A., Takagi, M., 2000. Gene regulation in response to overexpression of cytochrome P450 and proliferation of the endoplasmic reticulum in *Saccharomyces cerevisiae*. Biosci. Biotechnol. Biochem., 64(9), 1930–1936. 10.1271/bbb.64.1930

