## Supplemental file for "Optimizing cytochrome P450 activity for quillaic acid biosynthesis in *Saccharomyces cerevisiae*"

**Supplementary Materials**

**Supplementary Figures**

| Supplementary Figure S1 | The growth dynamics of yQA1 and yQA2 in 24-well plates monitored by Growth Profiler |
| --- | --- |
| Supplementary Figure S2 | MS spectra of 16α-hydroxy hederagenin produced by yQA2 |
| Supplementary Figure S3 | Venn diagram of proteins with significant change in different strains (yQA2 vs. yQA1) |
| Supplementary Figure S4 | Specific titres of target triterpenoids in yeasts expressing 1-3 copies of *P2* and *Q7* by GAL2 promoters |
| Supplementary Figure S5 | The specific titres (a) and distribution (b) of triterpenoids in yeasts expressing different CPRs |
| Supplementary Figure S6 | Fed-batch fermentation of strain yQA14 |

**Supplementary Tables**

| Supplementary Table S1 | Genes used in this work |
| --- | --- |
| Supplementary Table S2 | Plasmids used in this work |
| Supplementary Table S3 | *Saccharomyces cerevisiae* strains used in this work |
| Supplementary Table S4 | Primers used in this work |
| Supplementary Table S5 | NMR data for isolated oleanolic aldehyde |
| Supplementary Table S6 | NMR data for isolated 16α-hydroxy hederagenin |

**Supplementary Discussion**

Global proteomic changes and ergosterol pathway regulation

**Supplementary Figures**
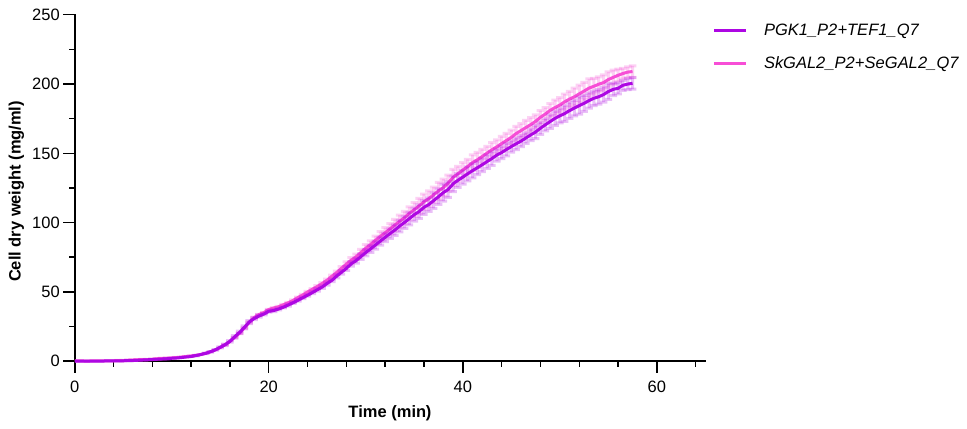

**Fig. S1 The growth dynamics of yQA1 and yQA2 in 24-well plates monitored by Growth Profiler**

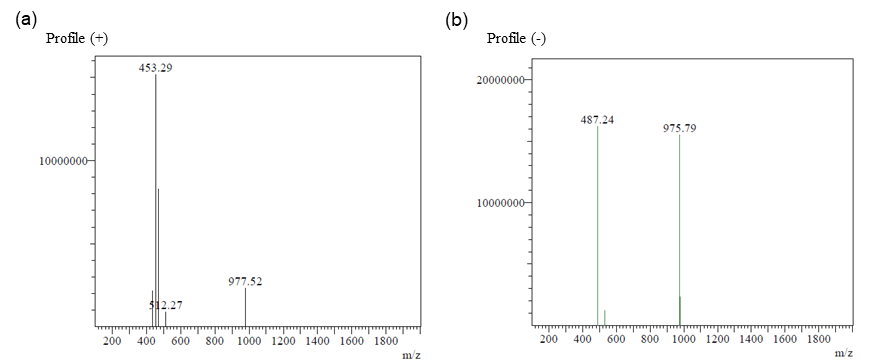

**Fig. S2 MS spectra of 16α-hydroxy hederagenin produced by yQA2.** (a) Profile(+) (Positive Ion Mode); (b) Profile(-) (Negative Ion Mode).

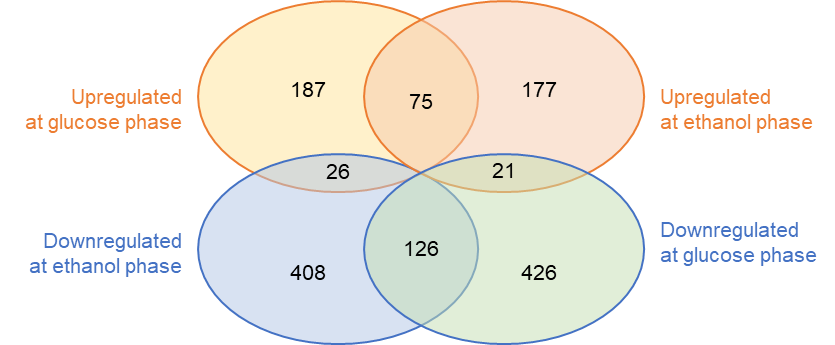

**Fig. S3 Venn diagram of proteins with significant change in different strains (yQA2 vs. yQA1)**

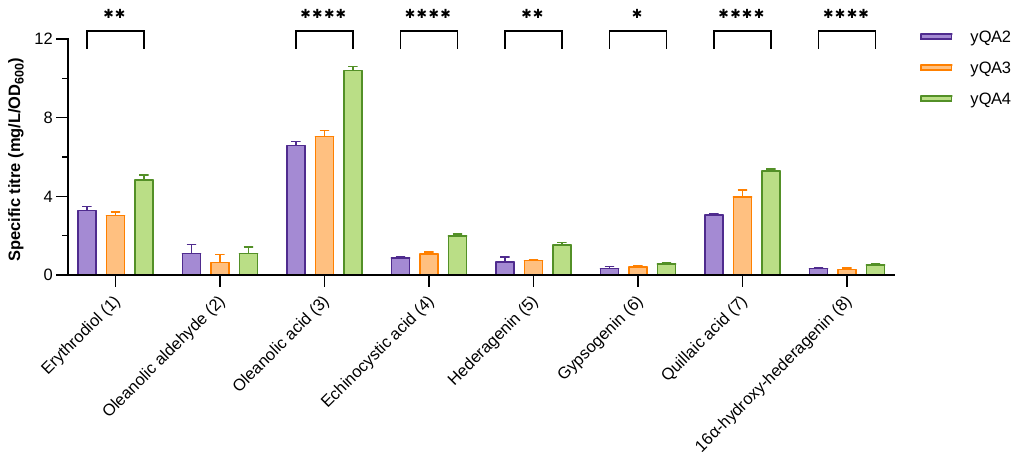

**Fig. S4 Specific titres of target triterpenoids in yeasts expressing 1-3 copies of *P2* and *Q7* by GAL2 promoters**

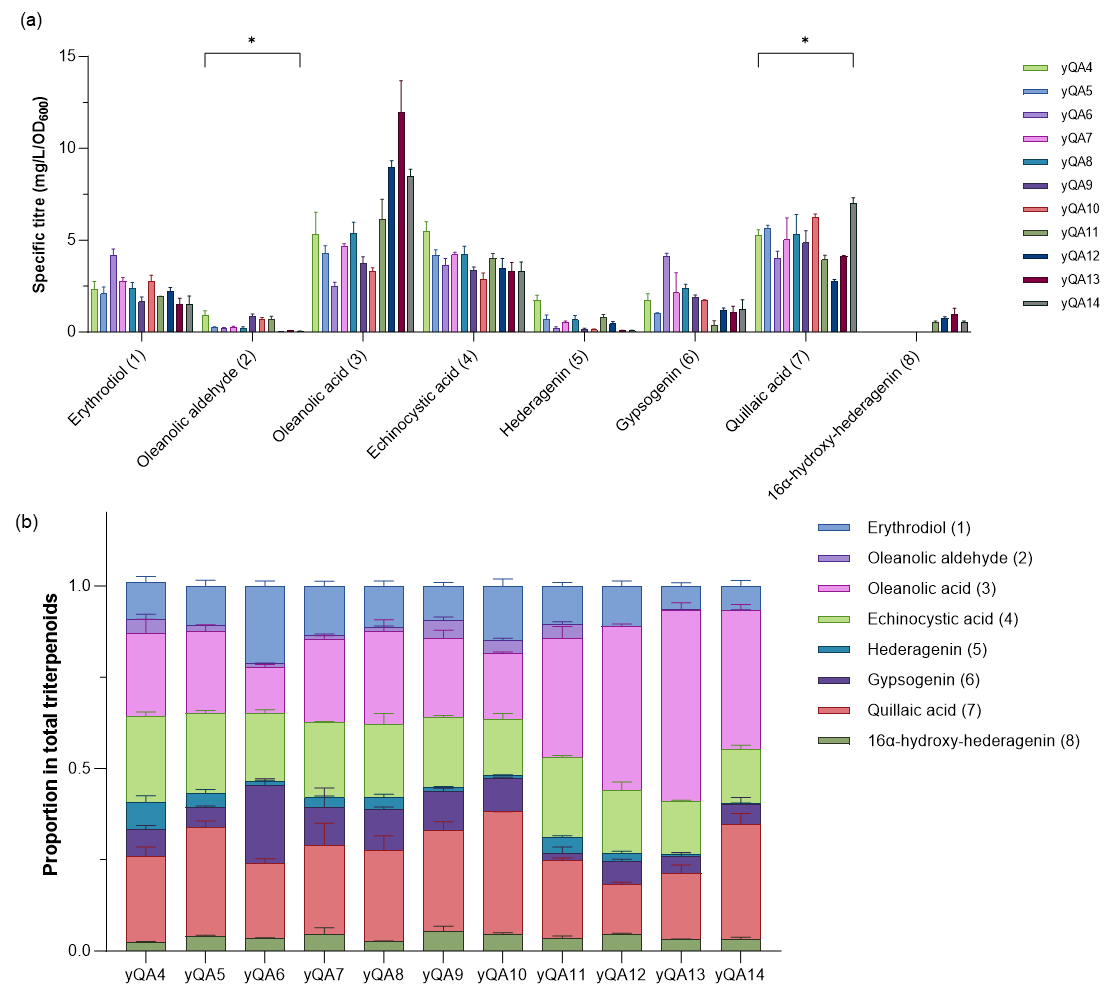

**Fig. S5 The specific titres (a) and distribution (b) of triterpenoids in yeasts expressing different CPRs**

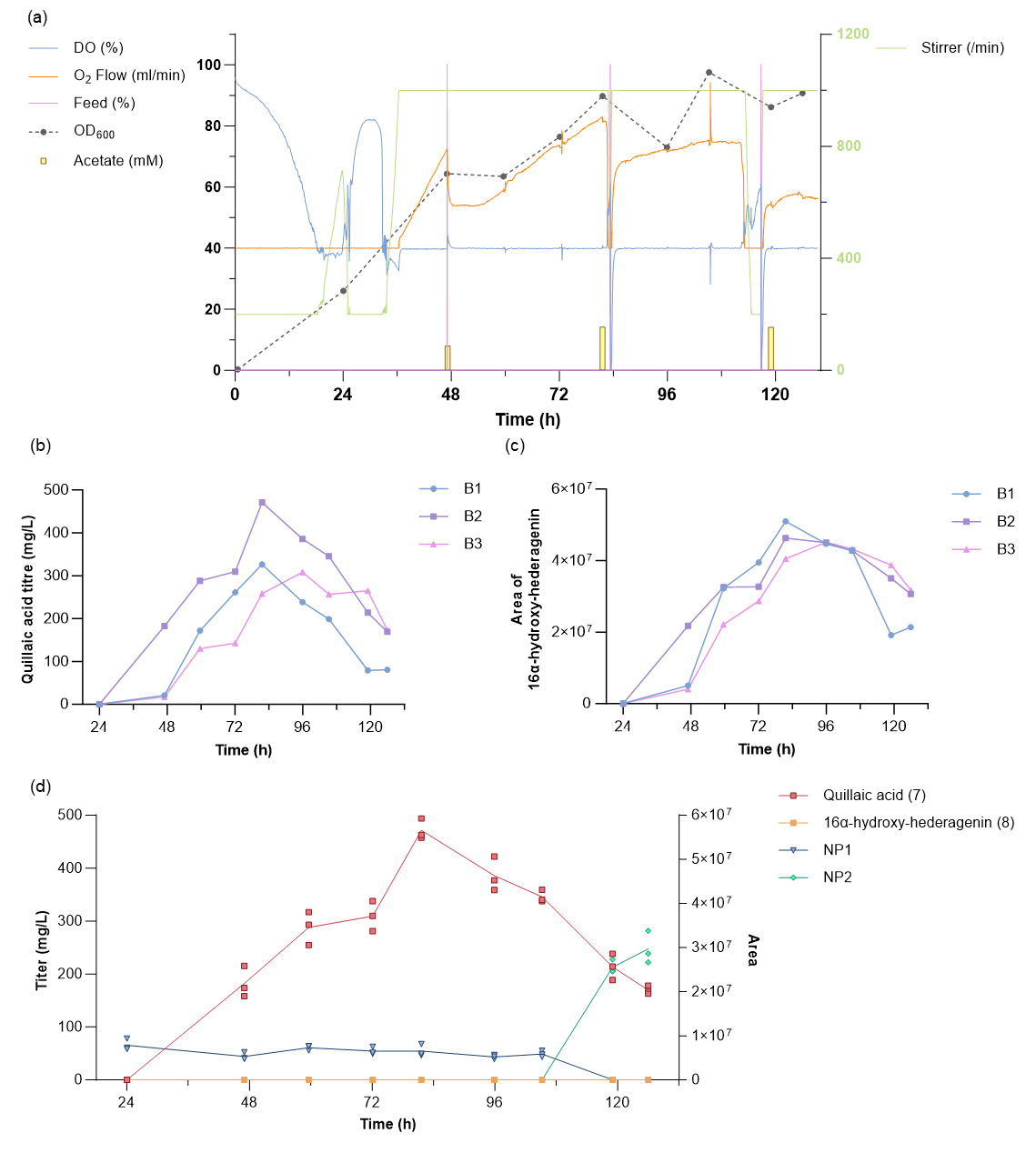

**Fig. S6 Fed-batch fermentation of strain yQA14.** (a) Key parameters recorded during fed-batch fermentation in BR2, along with the OD_600_ value over time. (b) Quillaic acid titre and (c) the area of 16α-hydroxy-hederagenin from the yeast during fed-batch fermentation in all the three bioreactors (d) Quillaic acid titre and peak area of the unidentified peaks from the yeast extract during fed-batch fermentation in bioreactors.

**Supplementary Tables**

**Table S1 Genes used in this work.**

| **Gene** | **Sequence (5’-3’)** |
| --- | --- |
| QsCYP714-7 (Q7) | MWFTVGLVLVFALFIRLYSSLWLKPRATRIKLSNQGIKGPKPAFLLGNVAEMRRFQSKLPKSELKQGQVSHDWASKSLFPFFSLWSQKYGNTFVFSLGNIQVLYVSDHELVKEINQNTSLDLGKPKYLQKERGPLLGQGILTSNGQLWAYQRKIMTPELYKEKIKGMCELMVESVAWLVEEWGTKIQAEGGAADIRIDEDLRSFSGDVISKACFGSCYAGGREIFLRLRALQHQIASKALLMGFPGLKYLPIKSNREIWRLEKEIFQLIMKLAEDRKKEQHERDLLQIIIEGAKSSDLSSEAMAKFIVDNCKNVYLAGHETTAMSAGWTLLLLANHPEWQARVRDEILQVTEGRNPDFDMLHKMKLLTMVIQEALRLYPTVIFMSREALEDINVGNIQVPKGVNIWIPVVNLQRDTTVWGADANEFNPERFANGVNNSCKVPQLYLPFGAGPRICPGINLAMTEIKILLCILLTKFSFSVSPNYRHSPVFKLVLEPENGINVIMKKL* |
| QsCYP716-201209 (Q9) | MIYNNDSNDNELVISSVQQPSMDPFFIFGLLLLALFLSVSFLLYLSRRAYASLPNPPPGKLGFPVVGESLEFLSTRRKGVPEKFVFDRMAKYCRDVFKTSILGATTAVMCGTAGNKFLFSNEKKHVTGWWPKSVELIFPTSLEKSSNEESIMMKQFLPNFLKPEPLQKYIPVMDIITQRHFNTSWEGRNVVKVFPTAAEFTTLLACRVFLSVEDPIEVAKISEPFEILAAGFLSIPINLPGTKLNKAVKAADQIRDAIVQILKRRRVEIAENKANGMQDIASMLLTTPTNAGFYMTEAHISEKILGMIVGGRDTASTVITFIIKYLAENPEIYNKVYEEQMEVVKSKKPGELLNWEDVQKMKYSWCVACEAMRLAPPVQGGFKVAINDFVYSGFNIRKGWKLYWSAIATHMNPEYFPEPEKFNPSRFEGKGPVPYSFVPFGGGPRMCPGKEYSRLETLVFMHHLVTRYNWEKVYPTEKITVDPMPFPVNGLPIRLIPHKHQ* |
| MtCYP72A68 (M8) | MELSWETKSAIILITVTFGLVYAWRVLNWMWLKPKKIEKLLREQGLQGNPYRLLLGDAKDYFVMQKKVQSKPMNLSDDIAPRVAPYIHHAVQTHGKKSFIWFGMKPWVILNEPEQIREVFNKMSEFPKVQYKFMKLITRGLVKLEGEKWSKHRRIINPAFHMEKLKIMTPTFLKSCNDLISNWEKMLSSNGSCEMDVWPSLQSLTSDVIARSSFGSSYEEGRKVFQLQIEQGELIMKNLMKSLIPLWRFLPTADHRKINENEKQIETTLKNIINKREKAIKAGEATENDLLGLLLESNHREIKEHGNVKNMGLSLEEVVGECRLFHVAGQETTSDLLVWTMVLLSRYPDWQERARKEVLEIFGNEKPDFDGLNKLKIMAMILYEVLRLYPPVTGVARKVENDIKLGDLTLYAGMEVYMPIVLIHHDCELWGDDAKIFNPERFSGGISKATNGRFSYFPFGAGPRICIGQNFSLLEAKMAMALILKNFSFELSQTYAHAPSVVLSVQPQHGAHVILRKIKT* |
| MlCYP87D16 (M6) | MWVVGLIGVAVVTILITQYVYKWRNPKTVGVLPPGSMGLPLIGETLQLLSRNPSLDLHPFIKSRIQRYGQIFATNIVGRPIIVTADPQLNNYLFQQEGRAVELWYLDSFQKLFNLEGANRPNAVGHIHKYVRSVYLSLFGVESLKTKLLADIEKTVRKNLIGGTTKGTFDAKHASANMVAVFAAKYLFGHDYEKSKEDVGSIIDNFVQGLLAFPLNVPGTKFHKCMKDKKRLESMITNKLKERIADPNSGQGDFLDQAVKDLNSEFFITETFIVSVTMGALFATVESVSTAIGLAFKFFAEHPWVLDDLKAEHEAVLSKREDRNSPLTWDEYRSMTHTMHFINEVVRLGNVFPGILRKALKDIPYNGYTIPSGWTIMIVTSTLAMNPEIFKDPLAFNPKRWRDIDPETQTKNFMPFGGGTRQCAGAELAKAFFATFLHVLISEYSWKKVKGGSVARTPMLSFEDGIFIEVTKKNK* |
| PtCYP716A262 (P2) | MDPITLLSGLLVLAIVSLPTFFILYYSHPTKDGKSLPPGRMGWPFIGESYDFFAAGWKGKPESFIFNRLAKYASGNLNGQFKTSLFGNKAVVVAGAAANKLLFSNEKKLVTMWWPPSIDKAFPSTAQLSANEEALLMRKFFPSFLIRREALQRYIPIMDDCTRRHFSTGAWGPSDRIEAFSVTQDYTFWVACRVFMSIDAQEDPATVDSLFKHFNVLKAGIYSMHIDLPWTNFHHAMKASQAIRSAVEQIAKKRRAELAEGKAFPTQDILSYMLETPISPAEENKDGKAKYLNDADIGTKILGLLVGGHDTSSTVITFFFKFMAENPHVYEAIYKEQMEVAASKEAGELLNWDDLQKMKYSWCAICEVMRLTPPVQGAFRQAITDFTHNGYLIPKGWKIYWSTHSTHRSPECFPQPEKFDPTRFEGNGPPPFSFVPFGGGPRMCPGKEYARLQVLTFVHHIVTKFKWEQILPNEKIVVSPMPYPEKNLPLRMIARS* |
| PtCYP716A257 (P7) | MKEYIPYVATGIACIVILRWVLNMLKWLWIEPRRLEKCLRKQGLEGNSYKFLFGDMKEASKLRTEALAKPMPMRFNHDYFPRINPFVDQLLNKYGTNCFMWMGPVPTVHIGEPELVREAFNRMHEFQKPKTNPLSALLATGLVSYEGDKWAKHRRLINPAFHVEKLKLMIPAFRESIVEVVKEWEKRVPESGSAEIDVWPYLTSLTGDVISRAAFGSVYGDGKRIFELLSVQKELVLSLLKYSYIPGYTYLPTEGNKKMKEVNNEIQRLLENVIQNRKKAMEAGEAAKDDLLGLLMDSNYKESLLEGGGKNKKLIMSFQDLIDECKLFFLAGHETTAVLLAWTMILLSKHQDWQTRAREEVLATFGMREPTDYDALNRLKTVTMILNEVLRLYPPVISTNRKLFNGEAKIGNLIIPSGVGISLLTVQANRDPKVWGEDASEFRPDRFAEGLVKATKGNVAFFPFGWGPRICIGQNFALTESKMAVAMILQRFTFDFSPTYTHAPSGLITLNPQYGAPLMFRKR* |
| MTR1 | MTSSNSDLVRTIESVLGVSLGDSVSDSVVLIVTTSAAVIIGLLVFLWKKSSDRSKELKPVIVPKSLVKEEDDDADIADGKTKVTVFFGTQTGTAEGFAKALAEEIKARYEKAFVKVVDMDDYAADDDQYEEKLKKETLAFFMLATYGDGEPTDNAARFYKWFTEGKDERGTWLQQLTYGVFGLGNRQYEHFNKIGKVVDDDLSEQGAKRLVPLGMGDDDQSIEDDFNAWKESLWPELDQLLRDEDDVNTVSTPYTAAISEYRVVFHDPTVTPSYENHFNAANGGAVFDIHHPCRANVAVRRELHKPQSDRSCIHLEFDVSGTGVTYETGDHVGVYADNCDETVKEAGKLLGQDLDLLFSLHTDNEDGTSLGGSLLPPFPGPCTVRTALARYADLLNPPRKAALIALAAHASEPSEAERLKFLSSPQGKDEYSKWVVGSHRTLLEVMADFPSAKPPLGVFFAAIAPRLQPRYYSISSSPRFAPQRVHVTCALVEGPTPTGRIHKGVCSTWMKNAIPSEESRDCSWAPIFIRPSNFKLPADPSIPIIMVGPGTGLAPFRGFLQERFALKEDGVQLGPALLFFGCRNRQMDFIYEEELNNFVEQGSLSELIVAFSREGPEKEYVQHKMMDKASYFWSLISQGGYLYVCGDAKGMARDVHRTLHTIVQQQENADSSKAEATVKKLQMDGRYLRDVW* |
| ATR2 | MSSSSSSSTSMIDLMAAIIKGEPVIVSDPANASAYESVAAELSSMLIENRQFAMIVTTSIAVLIGCIVMLVWRRSGSGNSKRVEPLKPLVIKPREEEIDDGRKKVTIFFGTQTGTAEGFAKALGEEAKARYEKTRFKIVDLDDYAADDDEYEEKLKKEDVAFFFLATYGDGEPTDNAARFYKWFTEGNDRGEWLKNLKYGVFGLGNRQYEHFNKVAKVVDDILVEQGAQRLVQVGLGDDDQCIEDDFTAWREALWPELDTILREEGDTAVATPYTAAVLEYRVSIHDSEDAKFNDINMANGNGYTVFDAQHPYKANVAVKRELHTPESDRSCIHLEFDIAGSGLTYETGDHVGVLCDNLSETVDEALRLLDMSPDTYFSLHAEKEDGTPISSSLPPPFPPCNLRTALTRYACLLSSPKKSALVALAAHASDPTEAERLKHLASPAGKDEYSKWVVESQRSLLEVMAEFPSAKPPLGVFFAGVAPRLQPRFYSISSSPKIAETRIHVTCALVYEKMPTGRIHKGVCSTWMKNAVPYEKSENCSSAPIFVRQSNFKLPSDSKVPIIMIGPGTGLAPFRGFLQERLALVESGVELGPSVLFFGCRNRRMDFIYEEELQRFVESGALAELSVAFSREGPTKEYVQHKMMDKASDIWNMISQGAYLYVCGDAKGMARDVHRSLHTIAQEQGSMDSTKAEGFVKNLQTSGRYLRDVW* |
| VvCPR1 | MSSESNLINTVEAFLGVSLGSESAVLILTTTVAVILGLLIFVWRRSSDRGRDVKPLVVPKAVAIPAEEDEAEAAVGKTKVTVFFGTQTGTAEGFAKALAEEIKARYDKATIKVVDLDDYAVDDDQYEEKLKKEALAFFMLATYGDGEPTDNAARFYKWFTEGKEREAWLQQLTYGVFGLGNRQYEHFNKIAKVVDELLSEQGAKRLVPAGLGDDDQCIEDDFAAWRELLWPELDLLLRDEDDISAVSTPYTAVIPEYRVVIHDPIVTSCEDKFLNMANGNASFDIHHPCRVNVAVQRELHTLESDRSCIHLEFDTSGTGITYETGDHVGVYAENCDETVEEAGRLLGQPLDLLFSIHTDKDDGTSLGSSLPPTFPGPCTIRTALACYADLLNSPRKAALSALAAHAIEPGEAERLKFLASPQGKDEYSQWVVGSQRSLLEVMAEFPSAKPPLGVFFAAIAPRLQPRYYSISSSPRYASHRVHVTCALVYGPSPTGRIHKGVCSTWMKNAVSLEKSHNSSWAPIFIRPSNFKLPVDPLTPIIMVGPGTGLAPFRGFLQERLALKEDGVQLGPALLFFGCRNRRMDFIYEDELNNFVEQGILSELIVAFSREGPQKEYVQHKMMDRASYIWNIISQGGYLYVCGDAKGMAKDVHRTLHTIVQEQENVESSKAEAIVKKLHTDGRYLRDVW* |
| VvCPR2 | MQSSSVKVSPFDLMSAIIKGSMDQSNVSSESGGAAAMVLENREFIMILTTSIAVLIGCVVVLIWRRSGQKQSKTPEPPKPLIVKDLEVEVDDGKQKVTIFFGTQTGTAEGFAKALAEEAKARYEKAIFKVVDLDDYAGDDDEYEEKLKKETLAFFFLATYGDGEPTDNAARFYKWFAEGKERGEWLQNLKYGVFGLGNRQYEHFNKVAKVVDDIITEQGGKRIVPVGLGDDDQCIEDDFAAWRELLWPELDQLLRDEDDATTVSTPYTAAVLEYRVVFHDPEGASLQDKSWGSANGHTVHDAQHPCRANVAVRKELHTPASDRSCTHLEFDISGTGLTYETGDHVGVYCENLPETVEEAERLLGFSPDVYFSIHTEREDGTPLSGSSLSPPFPPCTLRTALTRYADVLSSPKKSALVALAAHASDPSEADRLKYLASPSGKDEYAQWVVASQRSLLEIMAEFPSAKPPLGVFFAAVAPRLQPRYYSISSSPKMVPSRIHVTCALVCDKMPTGRIHKGICSTWMKYAVPLEESQDCSWAPIFVRQSNFKLPADTSVPIIMIGPGTGLAPFRGFLQERFALKEAGAELGSSILFFGCRNRKMDYIYEDELNGFVESGALSELIVAFSREGPTKEYVQHKMMEKASDIWNVISQGGYIYVCGDAKGMARDVHRTLHTILQEQGSLDSSKAESMVKNLQMTGRYLRDVW* |

**Table S2 Plasmids used in this work.**

| **Plasmid Name** | **Information** | **Backbone** | **Source** |
| --- | --- | --- | --- |
| pCfB2899(X-2 MarkerFree) | An integrative vector designed to target the genomic region ChrX: 194944..195980, with Amp marker |  | EasyClone-Marker Free kit |
| pCfB2904(XI-3 MarkerFree) | An integrative vector designed to target the genomic region ChrXI: 93378..94567, with Amp marker |  | EasyClone-Marker Free kit |
| pCfB3037(XI-5 MarkerFree) | An integrative vector designed to target the genomic region ChrXI: 11779..118967, with Amp marker |  | EasyClone-Marker Free kit |
| pCfB3040(XII-4 MarkerFree) | An integrative vector designed to target the genomic region ChrXII: 830227..831248, with Amp marker |  | EasyClone-Marker Free kit |
| pCfB2909(XII-5 MarkerFree) | An integrative vector designed to target the genomic region ChrXII: 839226..840357, with Amp marker |  | EasyClone-Marker Free kit |
| pCfB3020(gRNA X-2) | A gRNA helper vector designed to target the genomic region ChrX: 194944–195980, with Amp marker |  | EasyClone-Marker Free kit |
| pCfB3045(gRNA XI-3) | A gRNA helper vector designed to target the genomic region ChrXI: 93378..94567, with Amp marker |  | EasyClone-Marker Free kit |
| pCfB3046(gRNA XI-5) | A gRNA helper vector designed to target the genomic region ChrXI: 11779..118967, with Amp marker |  | EasyClone-Marker Free kit |
| pCfB3049(gRNA XII-4) | A gRNA helper vector designed to target the genomic region ChrXII: 830227..831248, with Amp marker |  | EasyClone-Marker Free kit |
| pCfB3050(gRNA XII-5) | A gRNA helper vector designed to target the genomic region ChrXII: 839226..840357, with Amp marker |  | EasyClone-Marker Free kit |
| pCfB3044(gRNA XI-2) | A gRNA helper vector designed to target the genomic region ChrXI: 91575..92913, with Amp marker |  | EasyClone-Marker Free kit |
| pYCYP1.1 | PGK1p>Q9 | pCfB3040 | This study |
| pYCYP1.2 | PGK1p>M6 | pCfB3040 | This study |
| pYCYP1.3 | PGK1p>P2 | pCfB3040 | This study |
| pYCYP1.4 | TEF1p>Q7 | pCfB3037 | This study |
| pYCYP1.5 | TEF1p>M8 | pCfB3037 | This study |
| pYCYP1.6 | TEF1p>P7 | pCfB3037 | This study |
| pYCYP2.1 | PGK1p>P7, TEF1p>Q7 | pCfB3040 | This study |
| pYCYP2.2 | SkGAL2p>P7, SeGAL2p>Q7 | pCfB3040 | This study |
| pYCYP2.3 | SkGAL2p>P7, SeGAL2p>Q7 | pCfB2899 | This study |
| pYCYP2.4 | SkGAL2p>P7, SeGAL2p>Q7 | pCfB3037 | This study |
| pYCPR1 | TEF1p>MTR1 | pCfB2904 | This study |
| pYCPR2 | TEF1p>ATR2 | pCfB2904 | This study |
| pYCPR3 | TEF1p>VvCPR1 | pCfB2904 | This study |
| pYCPR4 | TEF1p>VvCPR2 | pCfB2904 | This study |
| pYCPR5 | SkGAL2p>MTR1 | pCfB2909 | This study |
| pYCPR6 | SkGAL2p>VvCPR2 | pCfB2904 | This study |
| pYCPR7 | TEF1p>VvCPR2 | pCfB2909 | This study |
| pYCPR8 | SkGAL2p>VvCPR2 | pCfB2909 | This study |
| pYgal80g | A gRNA helper vector for GAL80 knock-out | pCfB3044 | This study |
| pYncp1g | A gRNA helper vector for NCP1 knock-out | pCfB3044 | This study |

**Table S3 *Saccharomyces cerevisiae* strains used in this work.**

| **Strain Name** | **Properties** | **Parent Strain** | **Source** |
| --- | --- | --- | --- |
| rOA2 | CEN.PK 102-5B, tHMG1, ATR2, are2::KanMX; single-copy integration of ADH1t<tHMG1<TEF1p-PGK1p>MTR>CYC1t cassettes and KlLEU2 marker gene into chromosome X; replacement of the native ERG7 gene with the frame-shift ERG7 mutant; CYC1t<AaBAS<PGK1p-TEF1p>CYP716A15>ADH1t cassettes and URA3 marker gene, CYC1t<AaBAS<PGK1p-TEF1p>CYP716A15>ADH1t cassettes and HIS3 marker gene sequentially integrated into chromosome XII |  | Lab stock |
| yQ9 | Integrative plasmid pYCYP1.1; PGK1p>Q9>CYC1t cassettes integrated into chromosome XII-4 | rOA2 | This study |
| yM6 | Integrative plasmid pYCYP1.2; PGK1p>M6>CYC1t cassettes integrated into chromosome XII-4 | rOA2 | This study |
| yP2 | Integrative plasmid pYCYP1.3; PGK1p>P2>CYC1t cassettes integrated into chromosome XII-4 | rOA2 | This study |
| yQ7 | Integrative plasmid pYCYP1.4; TEF1p>Q7>ADH1t cassettes integrated into chromosome XI-5 | rOA2 | This study |
| yM8 | Integrative plasmid pYCYP1.5; TEF1p>M8>ADH1t cassettes integrated into chromosome XI-5 | rOA2 | This study |
| yP7 | Integrative plasmid pYCYP1.6; TEF1p>P7>ADH1t cassettes integrated into chromosome XI-5 | rOA2 | This study |
| yQA1 | Integrative plasmid pYCYP2.1; CYC1t<P2<PGK1p-TEF1p>Q7>ADH1t cassettes integrated into chromosome XII-4 | rOA2 | This study |
| yQA2 | Integrative plasmid pYCYP2.2; CYC1t<P2<SkGAL2p-SeGAL2p>Q7>ADH1t cassettes integrated into chromosome XII-4; complete deletion of GAL80 | rOA2 | This study |
| yQA3 | Integrative plasmid pYCYP2.3; CYC1t<P2<SkGAL2p-SeGAL2p>Q7>ADH1t cassettes integrated into chromosome X-2 | yQA2 | This study |
| yQA4 | Integrative plasmid pYCYP2.4; CYC1t<P2<SkGAL2p-SeGAL2p>Q7>ADH1t cassettes integrated into chromosome XI-5 | yQA3 | This study |
| yQA5 | Integrative plasmid pYCPR1; TEF1p>MTR1>CYC1t cassettes integrated into chromosome XI-3 | yQA4 | This study |
| yQA6 | Integrative plasmid pYCPR2; TEF1p>ATR2>CYC1t cassettes integrated into chromosome XI-3 | yQA4 | This study |
| yQA7 | Integrative plasmid pYCPR3; TEF1p>VvCPR1>CYC1t cassettes integrated into chromosome XI-3 | yQA4 | This study |
| yQA8 | Integrative plasmid pYCPR4; TEF1p>VvCPR2>CYC1t cassettes integrated into chromosome XI-3 | yQA4 | This study |
| yQA9 | Integrative plasmid pYCPR5; SkGAL2p>MTR1>CYC1t cassettes integrated into chromosome XI-3 | yQA4 | This study |
| yQA10 | Integrative plasmid pYCPR6; SkGAL2p>VvCPR2>CYC1t cassettes integrated into chromosome XI-3 | yQA4 | This study |
| yQA11 | Integrative plasmid pYCPR7; TEF1p>VvCPR2>CYC1t cassettes integrated into chromosome XII-5 | yQA5 | This study |
| yQA12 | Integrative plasmid pYCPR7; TEF1p>VvCPR2>CYC1t cassettes integrated into chromosome XII-5 | yQA8 | This study |
| yQA13 | Integrative plasmid pYCPR8; SkGAL2p>VvCPR2>CYC1t cassettes integrated into chromosome XII-5; complete deletion of NCP1 | yQA10 | This study |
| yQA14 | Integrative plasmid pYCPR8; SkGAL2p>VvCPR2>CYC1t cassettes integrated into chromosome XII-5; complete deletion of NCP1 | yQA10 | This study |

**Table S4 Primers used in this work. Restriction enzyme sites are underlined.**

| **Primer Name** | **Sequence (5’-3’)** |
| --- | --- |
| PGK1-F | CGTATCTACCAACGGAATGCGTGAATTCGGCCGGCCTGGAAGTACCTTC |
| PGK1-Q9-R | CGTTCGAATCGTTGTTGTATATCATTTGTTTTATATTTGTTGTAAAAAGTAG |
| Q9-PGK1-F | CAACAAATATAAAACAAATGATATACAACAACGATTCGAACGACAATGAGCTAG |
| Q9-R | CTCCTTCCTTTTCGGTTAGAGCGGATGAATGCACGCGATCGCTTACTGGTGTTTATGC |
| PGK1-M6-R | TCCAATTAATCCAACTACCCACATTTGTTTTATATTTGTTGTAAAAAGTAGATAATTAC |
| M6-PGK1-F | TACAACAAATATAAAACAAATGTGGGTAGTTGGATTAATTGGAGTTGCAGTGGTGACG |
| M6-R | GTTAGAGCGGATGAATGCACGCGATCGCTTACTTGTTCTTTTTAGTCACTTCT |
| PGK1-P2-R | GCCACTTAACAAAGTGATTGGATCCATTTGTTTTATATTTGTTGTAAAAAGTAG |
| P2-PGK1-F | CAACAAATATAAAACAAATGGATCCAATCACTTTGTTAAGTGGCTTGCTTGTTTTGGC |
| P2-R | GGTTAGAGCGGATGAATGCACGCGTGATCTCGCAATCATTCTTAGG |
| TEF1-F | GGTTAGAGCGGATGAATGCACGCGTGATCTCGCAATCATTCTTAGG |
| TEF1-Q7-R | AAACCTACTGTAAACCACATTTGTAATTAAAACTTAGATTAGATTGCTATGCTTTCTTTC |
| Q7-TEF1-F | CAATCTAATCTAAGTTTTAATTACAAATGTGGTTTACAGTAGGTTTAGTTCTGGTTTTT |
| Q7-R | ACAACGTATCTACCAACGGAATGCGTAAGCTTTTATAGTTTTTTCATGATAACGTTAATC |
| TEF1-M8-R | GACTTTGTCTCCCAGGACAACTCCATTTGTAATTAAAACTTAGATTAGATTGCTATGC |
| M8-TEF1-F | ATCTAAGTTTTAATTACAAATGGAGTTGTCCTGGGAGACAAAGTCAGCCATTAT |
| M8-R | CAACGTATCTACCAACGGAATGCGTAAGCTTTTAGGTCTTAATCTTTCTAAGAATAAC |
| TEF1-P7-R | GCAACGTATGGGATATATTCCTTCATTTGTAATTAAAACTTAGATTAGATTGC |
| P7-TEF1-F | CTAAGTTTTAATTACAAATGAAGGAATATATCCCATACGTTGCAACAGGTATTGC |
| P7-R | CTACCAACGGAATGCGTGCGATACGTTTTCTGAACATCAGGGGG |
| TEF1-PGK1-F | TACTTCCAGGCCGGCCCATATGGCACACACCATAGCTTCAAAATGTTTCTACTCCTT |
| SkGAL2-F | GTATCTACCAACGGAATGCGTGCGATCGCTAAACCAATTTTATTTGAACTTGCCCCG |
| SkGAL2-P2-F | CAAAGTGATTGGATCCATGGTACCTGTAAAAAACTTTTTTTATTATACTATTTTC |
| P2-SkGAL2-F | GTTTTTTACAGGTACCATGGATCCAATCACTTTGTTAAGTGGCTTGCTTG |
| SeGAL2-SkGAL2-F | AATAAAATTGGTTTAGCGATCGCCCACAGAGAACAGGAGATTACAAGATCCAG |
| SeGAL2-Q7-F | CCTACTGTAAACCACATGAATTCTGTAAATGTGTGTATATATTATATTATAGTATTG |
| Q7-SeGAL2-F | CACACATTTACAGAATTCATGTGGTTTACAGTAGGTTTAGTTCTGGTTTTTG |
| TEF1-MTR1-R | CTATTGGAGGAAGTCATGTCGACTTGTAATTAAAACTTAGATTAGATTGCTATG |
| MTR1-TEF1-F | CTAAGTTTTAATTACAAGTCGACATGACTTCCTCCAATAGTGATCTAGTGCG |
| MTR1-R | GGTTAGAGCGGATGAATGCACGCGTCACCAAACGTCTCTCAAGTAC |
| TEF1-ATR2-R | GAAGAAGAGGAGGACATGTCGACTTGTAATTAAAACTTAGATTAGATTGCTATGC |
| ATR2-TEF1-F | CTAAGTTTTAATTACAAGTCGACATGTCCTCCTCTTCTTCATCATCCACCTCTATG |
| ATR2-R | CTTTTCGGTTAGAGCGGATGAATGCACGCGCGCGATTCACCAGACATCTCTCAAGTATC |
| TEF1-VvCPR1-R | GTTAGATTCAGAAGACATGTCGACTTGTAATTAAAACTTAGATTAGATTGCTATGC |
| VvCPR1-TEF1-F | CTAATCTAAGTTTTAATTACAAGTCGACATGTCTTCTGAATCTAACTTGATCAACACTG |
| VvCPR1-R | GGATGAATGCACGCGGCGATCGCTTACCAGACATCTCTTAGATATC |
| TEF1-VvCPR2-R | CAGAAGAAGATTGCATGTCGACTTGTAATTAAAACTTAGATTAGATTGCTATG |
| VvCPR2-TEF1-F | CTAAGTTTTAATTACAAGTCGACATGCAATCTTCTTCTGTTAAGGTTAGTCC |
| VvCPR2-R | GGTTAGAGCGGATGAATGCACGCGTTACCAGACATCTCTTAGATATCTACC |
| SkGAL2-MTR1-R | CTAGATCACTATTGGAGGAAGTCATGTCGACTGTAAAAAACTTTTTTTATTATAC |
| MTR1-SkGAL2-F | AAAAGTTTTTTACAGTCGACATGACTTCCTCCAATAGTGATCTAGTGCGTACC |
| SkGAL2-VvCPR2-R | GGACTAACCTTAACAGAAGAAGATTGCATGTCGACTGTAAAAAACTTTTTTTATTATAC |
| VvCPR2-SkGAL2-F | AAAAGTTTTTTACAGTCGACATGCAATCTTCTTCTGTTAAGGTTAGTCCATTCGATC |
| Upgal80-F | ATCGTGGAATTTATCCTGCT |
| Upgal80-R | AAGCACAGGGCAAGATGCTTGACGGGAGTGGAAAGAACGG |
| Downgal80-F | CCGTTCTTTCCACTCCCGTCAAGCATCTTGCCCTGTG |
| Downgal80-R | ATGATTCCATGCTACCTTCC |
| gRNAUp-F | ATCGGTGCGGGCCTCTTCGCTATTACGCCAGCTGGAATGCGTGCGATAGGGAACAAAAG |
| gRNAgal80Up-R | CTAGCTCTAAAACAATCGGTTACCACGCTCATCGATCATTTATCTTTCACTGCGGAG |
| gRNAgal80Down-F | GTGAAAGATAAATGATCGATGAGCGTGGTAACCGATTGTTTTAGAGCTAGAAATAG |
| gRNADown-R | CGCGCGTTGGCCGATTCATTAATGCAGCTGGAATGCACGCGATTAACTAATTACATGAC |
| Upncp1-F | CGCACTACCCATCTATATAGCTATGTATTCTATATCCACG |
| Upncp1-R | GAAGGCTTGATCAGTGGGCTGagATATGTGCGACAGTAGATCACTTGAAAAC |
| Downncp1-F | GTGATCTACTGTCGCACATATctCAGCCCACTGATCAAGCCTTCGGCGCGGTTG |
| Downncp1-R | GAAAGGAAATGGGCTATCGTTACCTCTGGTTCTCCATATTT |
| gRNAncp1Up-R | GCTCTAAAACCATGGGTACGGTGTCCCATTGATCATTTATCTTTCACTGCGGAGAAG |
| gRNAncp1Down-F | GAAAGATAAATGATCAATGGGACACCGTACCCATGGTTTTAGAGCTAGAAATAGCAAG |

**Table S5 NMR data for isolated oleanolic aldehyde**

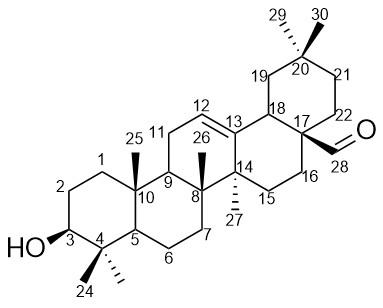

| 700 MHz, methanol-d_4_ + ~0.1% TFA | | |
| --- | --- | --- |
|  | **δ ^1^H (mult, *J*)** | **δ ^13^C** |
| 1 | 0.97-1.01 (ov)  1.62-1.66 (ov) | 39.8 |
| 2 | 1.61-1.65 (ov)  1.54- 1.58 (ov) | 27.8 |
| 3 | 3.15 (dd, 11.5, 4.6) | 79.6 |
| 4 | - | 39.8 |
| 5 | - | 56.7 |
| 6 | 1.42 (dd, 12.2, 3.2)  1.55-1.58 (ov) | 19.5 |
| 7 | 1.31 (dt, 12.8, 3.1)  1.49- 1.53 (ov) | 34.0 |
| 8 | - | 40.8 |
| 9 | 1.56-1.60 (ov) | 48.8^a^ |
| 10 | - | 38.1 |
| 11 | 1.89-1.95 (m, ov) | 24.5 |
| 12 | 5.41 (t, 3.67) | 124.5 |
| 13 | - | 144.4 |
| 14 | - | 42.8 |
| 15 | 1.10 (ddd, 14.0, 4.5, 2.7)  1.69 (ddd, 14.0, 14.0, 4.8) | 27.8 |
| 16 | 1.55- 1.59 (ov)  2.05 (ddd, 14.0, 14.0, 4.5) | 23.0 |
| 17 | - | 50.3 |
| 18 | 2.79 (dd, 13.8, 4.7) | 41.8 |
| 19 | 1.17-1.20 (ov)  1.76 (t, 13.7) | 46.8 |
| 20 | - | 31.6 |
| 21 | 1.38 (ddd, 13.6, 13.6, 4.1)  1.24 - 1.28 (m) | 34.1 |
| 22 | 1.49 (td, 13.6, 4.4)  1.17 - 1.21 (ov) | 28.8 |
| 23 | 0.78 (s)^b^ | 16.3^b^ |
| 24 | 0.97 (s)^b^ | 28.7^b^ |
| 25 | 0.95 (s) | 15.9 |
| 26 | 0.76 (s) | 17.7 |
| 27 | 1.17 (s) | 26.0 |
| 28 | 9.60 (s) | 209.2 |
| 29 | 0.93 (s)^c^ | 33.5^c^ |
| 30 | 0.94 (s)^c^ | 23.8^c^ |

s = singlet, m = multiplet, dd = doublet of doublets, dt = doublet of triplets, ddd = doublet of doublets of doublets, t = triplet, td = triplet of doublets, ov = overlapped signal

^a^ Signal is obscured by solvent. The chemical shift is taken from the HSQC spectrum.

^b^ Signal assignments may be reversed

^c^ Signal assignments may be reversed.

**Table S6 NMR data for isolated 16α-hydroxy hederagenin**

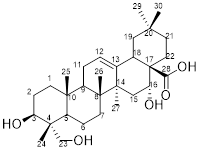

| 700 MHz, methanol-d_4_ + ~0.1% TFA | | |
| --- | --- | --- |
|  | **δ ^1^H (mult, *J*)** | **δ ^13^C** |
| 1 | 0.99 - 1.03 (ov)  1.63 - 1.66 (ov) | 39.6 |
| 2 | 1.58 - 1.62 (ov)  1.65 - 1.69 (ov) | 27.5 |
| 3 | 3.61 (dd, 11.7, 4.7) | 74.0 |
| 4 | - | 43.3 |
| 5 | 1.14 - 1.17 (m) | 49.0^a^ |
| 6 | 1.36 - 1.42 (ov)  1.44 - 1.48 (m) | 19.2 |
| 7 | 1.16 - 1.67 (ov)  1.27 - 1.33 (ov) | 33.8 |
| 8 | - | 40.6 |
| 9 | 1.68 - 1.73 (ov) | 48.2 |
| 10 | - | 37.9 |
| 11 | 1.84 - 1.89 (ov) | 24.5 |
| 12 | 5.29 (t, br, 3.6) | 124.7 |
| 13 | - | 146.5 |
| 14 | - | 42.8 |
| 15 | 1.33-1.36 (m)  1.86 (d, br) | 36.3 |
| 16 | 4.47 (s, br) | 75.5^b^ |
| 17 | - | Not obs |
| 18 | 3.01 (d, br, 13.2) | Not obs |
| 19 | 1.04 (d, br, 13.2)  2.30 (t, 13.2) | 47.9 |
| 20 | - | 31.4 |
| 21 | 1.14-1.18 (ov)  1.92-1.97 (ov) | 36.6 |
| 22 | 1.89-1.94 (ov)  1.76-1.81 (m) | 32.7 |
| 23 | 3.29 - 3.31 (ov)  3.52-3.55 (d, 10.9) | 67.4 |
| 24 | 0.71 (s) | 12.7 |
| 25 | 0.99 (s) | 16.4 |
| 26 | 0.81 (s) | 17.8 |
| 27 | 1.39 (s) | 27.5 |
| 28 | - | Not obs. |
| 29 | 0.88 (s)^c^ | 33.4^c^ |
| 30 | 0.97 (s)^c^ | 24.9^c^ |

s = singlet, m = multiplet, dd = doublet of doublets, t = triplet, br = broad signal, ov = overlapped signal, Not obs = signal not observed.

^a^ Signal is obscured by solvent. The chemical shift is taken from the HSQC spectrum.

^b^ chemical shift is taken from the HSQC spectrum

^c^ Signal assignments may be reversed

**Supplementary Discussion**

**Global proteomic changes and ergosterol pathway regulation**

A global comparison between yQA2 and yQA1 (Fig. S3) identified 950 differentially expressed proteins across both growth phases, with 661 showing reduced abundance in yQA2.

During growth on ethanol, most proteins involved in ergosterol synthesis were repressed in yQA1 and yQA2 relative to glucose (Fig. 4), with stronger downregulation in yQA2 compared to yQA1 during the ethanol phase. Oxygen limitation during ethanol metabolism may have impaired ergosterol biosynthesis, known to be differentially expressed under hypoxic conditions. This is supported by the significantly lower expression of several enzymes, Erg11p, Erg25p and Erg5p, which require molecular oxygen as the electron acceptor (Jordá & Puig, 2020). Considering that P2 and Q7 were strongly overexpressed at this stage in yQA2, ER stress can also be a key factor affecting Erg protein abundance, particularly for those primarily localized to the ER, such as Erg2p and Erg5p (Kristan and Rižner, 2012). In contrast, Erg6p, which is mostly located in lipid particles (LPs) (Leber et al., 1998), was upregulated in yQA2. The upregulation of proteins implicated in ER membrane biogenesis is linked to the UPR, a cellular reaction to the accumulation of misfolded or unfolded proteins within the ER. The UPR drives the enlargement of the ER membrane to meet the increased demand for protein processing (Kim et al., 2008). Ire1p (inositol requiring endonuclease 1) plays a critical role in ER stress by initiating the expression of enzymes that promote ER membrane growth such as Ino1p (inositol-3-phosphate synthase) implicated in phospholipid biosynthesis, which was upregulated in yQA2 at ethanol phase. Ire1p also helps alleviate ER stress by inducing chaperones, resulting in increased capacity to process nascent proteins (Schuck et al., 2009). Expression of the Kar2p chaperone is activated through this Ire1p-Hac1p pathway (Wang and Sevier, 2016), and in yQA2, further suggesting that the ER stress response was activated due to CYP450 overexpression.

**References**

Jordá, T., Puig, S., 2020. Regulation of ergosterol biosynthesis in *Saccharomyces cerevisiae*. Genes, 11(7), 795. <https://doi.org/10.3390/genes11070795>

Kim, I., Xu, W., Reed, J. C., 2008. Cell death and endoplasmic reticulum stress: Disease relevance and therapeutic opportunities. Nat. Rev. Drug Discov., 7(12), 1013-1030. <https://doi.org/10.1038/nrd2755>

Kristan, K., Rižner, T. L., 2012. Steroid-transforming enzymes in fungi. J. Steroid Biochem. Mol. Biol., 129(1), 79-91. <https://doi.org/https://doi.org/10.1016/j.jsbmb.2011.08.012>

Leber, R., Landl, K., Zinser, E., Ahorn, H., Spök, A., Kohlwein, S. D., Turnowsky, F., Daum, G., 1998. Dual localization of squalene epoxidase, Erg1p, in yeast reflects a relationship between the endoplasmic reticulum and lipid particles. Mol. Biol. Cell, 9(2), 375-386. <https://doi.org/10.1091/mbc.9.2.375>

Schuck, S., Prinz, W. A., Thorn, K. S., Voss, C., Walter, P., 2009. Membrane expansion alleviates endoplasmic reticulum stress independently of the unfolded protein response. J. Cell Biol., 187(4), 525-536. <https://doi.org/10.1083/jcb.200907074>

Wang, J., Sevier, C. S., 2016. Formation and reversibility of bip protein cysteine oxidation facilitate cell survival during and post oxidative stress. J. Biol. Chem., 291(14), 7541-7557. <https://doi.org/10.1074/jbc.M115.694810>
